# Post-glacial expansion dynamics, not refugial isolation, shaped the genetic structure of a migratory bird, the yellow warbler

**DOI:** 10.1101/2021.05.10.443405

**Authors:** Eleanor F. Miller, Michela Leonardi, Zhe Xue, Robert Beyer, Mario Krapp, Marius Somveille, Gian Luigi Somma, Pierpaolo Maisano Delser, Andrea Manica

## Abstract

During the glacial periods of the Pleistocene, swathes of the Northern Hemisphere were covered by ice sheets, tundra, and permafrost, leaving large areas uninhabitable for temperate and boreal species. The glacial refugia paradigm proposes that, during glaciations, species living in the Northern Hemisphere were forced southwards, forming isolated populations that persisted in disjunct regions known as refugia. According to this hypothesis, as ice sheets retreated, species recolonised the continent from these glacial refugia, and the mixing of these lineages is responsible for modern patterns of genetic diversity. An alternative hypothesis is that complex genetic patterns could also arise simply from heterogenous post-glacial expansion dynamics, without separate refugia. Both mitochondrial and genomic data from the North American yellow warbler (*Setophaga petechia)* shows the presence of an eastern and western clade, a pattern often ascribed to the presence of two refugia. However, species distribution modelling (SDM) of the past range of this species fails to identify obvious refugia during the Last Glacial Maximum. Using a climate-informed spatial genetic modelling (CISGeM) framework, which allows us to integrate knowledge of past geographic ranges based on SDM, we reconstructed past population sizes, range expansions, and likely recolonisation dynamics of this species, generating spatially and temporally explicit demographic reconstructions. The model captures the empirical genetic structure despite including only a single, large glacial refugium. The observed contemporary population structure was generated during the expansion dynamics after the glaciation and is due to unbalanced rates of northward advance to the east and west linked to the melting of the icesheets. Thus, modern population structure in this species is consistent with expansion dynamics, and refugial isolation is not required to explain it, highlighting the importance of explicitly testing drivers of geographic structure.

**Significance statement:** Patterns of population differentiation in many species have often been attributed to the mixing of isolates from distinct refugia that formed during periods of glaciation, when range fragmentation was likely. By formally bringing together multiple lines of evidence, we demonstrate that the patterns of genetic diversity seen across the range of the yellow warbler (*Setophaga petechia*) were not the result of multiple isolated refugia. Instead, asymmetric expansion from a single cohesive range generated the observed patterns; the expansion’s asymmetry was due to the uneven melting of the icesheets over time. Thus, we demonstrate the importance of reconstructing species’ range dynamics when trying to explain patterns of genetic differentiation.

## Introduction

It has frequently been shown that seemingly continuously distributed populations in the Northern Hemisphere harbour geographic structure in their genetic diversity. Indeed, within North America, many widespread and migratory passerines exhibit clear differences in both migration patterns and genomic diversity between eastern and western populations e.g. (1–3). This pattern has been interpreted as the consequence of glaciations, during which species were forced southwards, forming isolated, insular populations that persisted in disjunct regions known as refugia (4,5). According to this narrative, as ice-sheets retreated, species recolonised the continent from these glacial refugia, and the subsequent mixing of these lineages is responsible for modern patterns of genetic diversity.

However, even though the cycles of expansion and contraction could have fragmented ranges, leading to multiple glacial refugia in some species, multiple glacial refugia have not been demonstrated for all species e.g. (6,7). Indeed, it is becoming clear that glaciations in North America might not have driven range fragmentation as ubiquitously as it has previously been assumed, e.g. (8,9).

What other processes might then have shaped the genetics of modern populations? Range expansions have been shown to have the potential to leave profound signatures in the genetic structure of metapopulations through repeated founder events (10). An extreme consequence of this process is gene surfing, when rare variants can become common through stochastic sampling during a founder event, and then be spread widely at high frequency during the subsequent expansion (11). An important role for the recolonization dynamics in shaping modern-day population structuring has been recently put forward for a trans-continentally distributed species, the painted-turtle, *Chrysemys picta* (12). Reid et al. (12) demonstrated that, for this species, genetic differentiation during range expansion and isolation-by-distance are more likely to have driven modern-day population diversity than isolation in allopatric refugia.

The difficulty in quantifying the role of the range change dynamics on the genetic structure of species is that the results are highly dependent on the detail of the dynamics. Whilst it is straightforward to build simple spatial models that represent a range expansion (e.g. using SPLATCHE (13)), capturing the spatial and temporal heterogeneities of the real process is challenging. A possible solution is to use climate informed spatial genetic models (CISGeMs), which use climate reconstructions to condition the local demography of individual populations within a geographically explicit environment. This approach has been used in the past to directly estimate demographic parameters that underpin the effect of climate for the dynamics of the out of Africa expansion of humans (14,15).

Here, we demonstrate how CISGeMs can also be used to explicitly integrate information on the possible past range of a species into population genetics models, thus allowing us to combine quantitatively different lines of evidence (as encouraged by (16)). Specifically, we use Species Distribution Models (SDMs) (17), which quantify the bioclimatic niche of a species based on its present-day distribution, and backcast its range according to climatic changes. It is thus possible to formally verify whether the geographic range changes inferred by SDM can be reconciled with the genetics. If so, it is also possible to explore which demographic parameters would be most supported based on Approximate Bayesian Computation by comparing predicted and observed genetic quantities (see Fig. 1).

**Figure 1.**
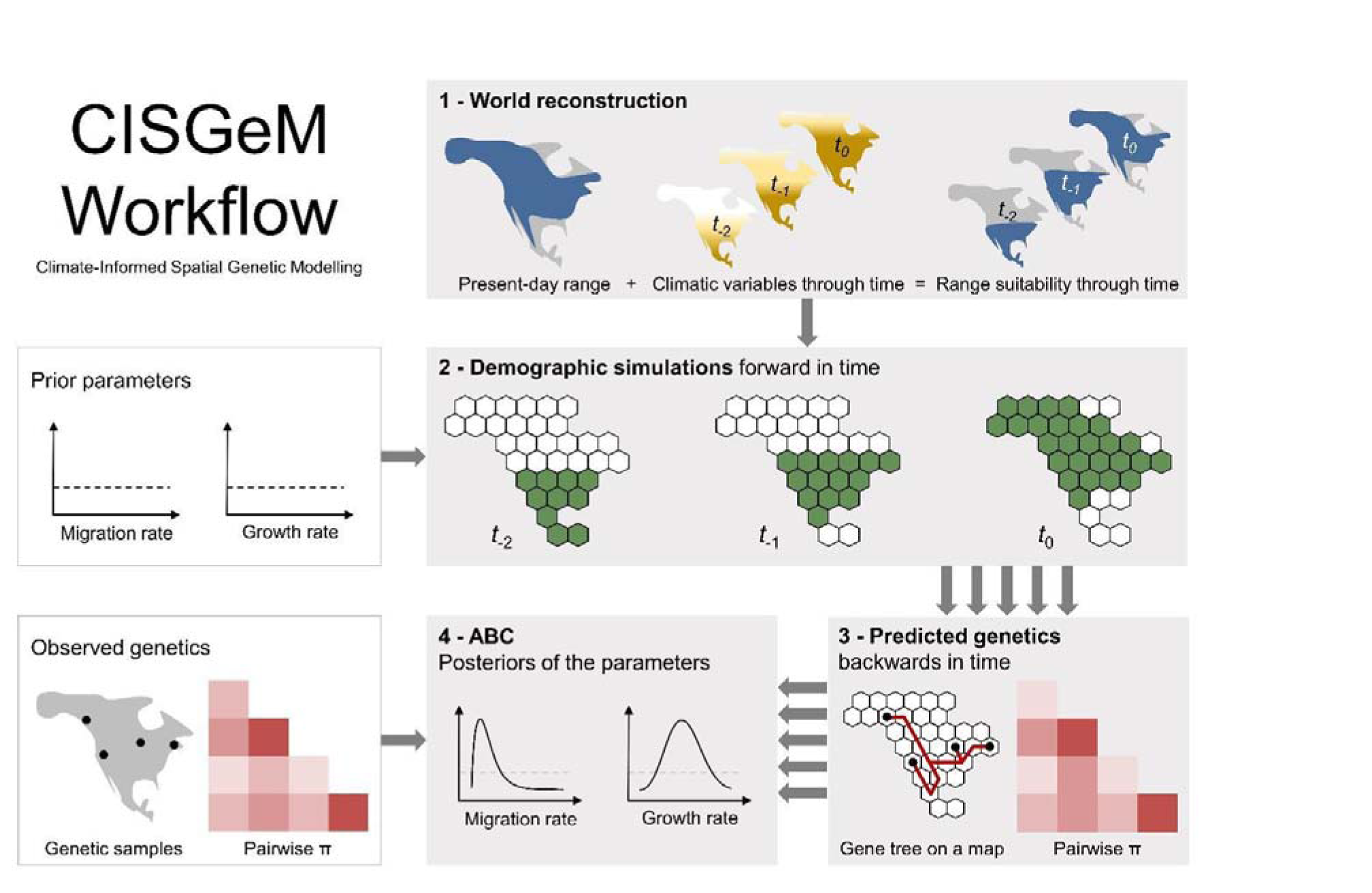
A schematic representation of the Climate-Informed Spatial Genetic Modelling (CISGeM) framework implemented in this paper.

In this paper, we use the CISGeM framework to explore the past range dynamics of the yellow warbler. This species is an abundant passerine species with a large continuous contemporary range and clear geographic population structuring, for which range-wide genomic data are available (18). Here we test whether today’s patterns of genetic structure in the North American yellow warbler (*Setophaga petechia*) can be best explained by recolonization from isolated glacial refugia, or if, more simply, heterogenous post-glacial expansion dynamics, without separate refugia, may have been sufficient to generate the patterns observed today. Firstly, we describe the genetic patterns that are found in the yellow warbler today from an empirical RAD-seq dataset. Then, we fit an SDM to reconstruct range changes through time. Finally, we use a CISGeM to explicitly integrate these two lines of evidence, test whether they are coherent, and to determine what were the drivers behind modern genomic patterns.

## Results

### Observed genetics

The 200 samples included in our study came from 21 sites across the modern breeding range of the North American yellow warbler. Sample sizes per site ranged from 6 to 20 individuals (see Materials and Methods). Analysis of the genetic structure (19) of the yellow warbler population revealed a clear longitudinal divide, with distinct East and West clusters that admix in the centre of the continent. This pattern is congruent with both the distribution of mitochondrial haplotypes (20) and patterns of migratory connectivity (21) in this species.

### Species Distribution Modelling for world reconstruction

We built a Species Distribution Model for yellow warblers based on modern data and projected back in time using paleoclimate reconstructions (step 1 in Fig. 1). The raw species occurrences data to define the present-day range was downloaded from the Global Biodiversity Information facility (GBIF, our data can be found at https://doi.org/10.15468/dl.jfkwcg) and totalled 1,573,147 data points. After filtering for coordinate accuracy, allowing an attributed error of 1km maximum, and filtering to only include points found within the BirdLife breeding and resident geographical ranges (22), we were left with 177,202 data points. As SDM works on presence/absence data and not frequencies, we retained only one presence per 0.5° grid cell, further refining this dataset down to 3,364 observations. With these observations we selected the four most informative, uncorrelated (threshold=0.7), bioclimatic variables to base our model on: Leaf Area Index (LAI), BIO7 (Temperature Annual Range), BIO8 (Mean Temperature of Wettest Quarter), BIO14 (Precipitation of Driest Month). Observations were further thinned based on a minimum distance between points of 70 km, leaving 1,188 presences; this procedure is used to correct for uneven sampling biases (23). We fitted SDMs to predict the probability of occurrence in each grid cell. We used an ensemble of four different algorithms: generalised linear models (GLM, (24)), generalized boosting method (GBM, (25)), generalised additive models (GAM, (26)), and random forest (27). Models were run performing spatial cross validation with 80% of the data used to train the algorithm and the remaining 20% to test it.

At present, the predicted potential distribution matches well the best range estimates for the species (Supplementary Fig. 1). Based on paleoclimate and vegetation reconstruction (see Materials and Methods), range projections from the present day back to 50 thousand years ago suggest that the distribution of habitat suitable for the yellow warbler expanded and contracted, to various degrees, multiple times. The potential range underwent a substantial contraction into the south of the continent at the peak of the Last Glacial Maximum (LGM), ∼21kya, before beginning to re-expand (Supplementary Fig. 2, Fig. 2 B-C), but the range never separated into distinct eastern and western refugia. Following the LGM, the retreat of the Cordillerian Ice Sheet was asymmetric: in the west, the ice started to retreat at about 18kya (28) with the opening of a corridor that progressively expanded to the higher latitudes, whereas the eastern and central part of the Laurentide Ice Sheet began retreating much later (29). By 13kya this deglaciated terrain became habitable for yellow warblers according to our SDM (Supplementary Fig. 2. C). From then on, as the ice sheets retreated further, habitat to the east of the continent and in the central area deglaciated, becoming increasingly viable (Supplementary Fig. 2. D-F).

**Figure 2.**
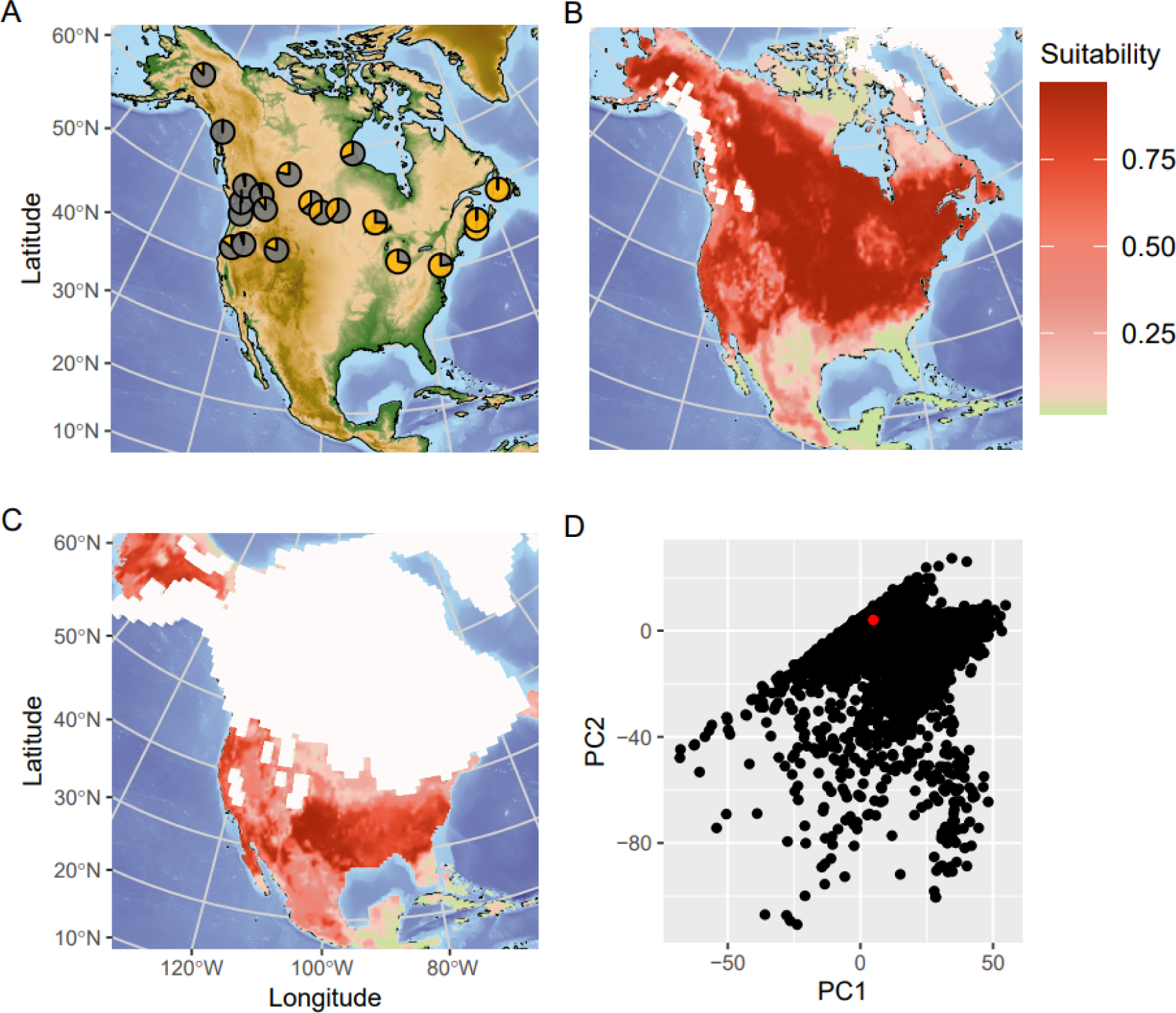
A) Genetic clustering (*K* = 2) results for all 21 populations in our study. Populations in the East and West of the continent are mostly uniform, whilst those in the central part are a mix of the two groups. This pattern is congruent with the distribution of mtDNA haplotypes (20). B) The potential range for the North American yellow warbler in the present day, reconstructed with an SDM. C) The potential range for the North American yellow warbler at 21kya, around the peak of the ice age. D) Pairwise plots of the first 2 PCs of summary statistics distributions from a Monte-Carlo sweep using CISGeM demonstrating that the model is able to capture the observed genetic patterns. Red dot represents observed value, simulated values are in black.

### Climate Informed Spatial Genetic Model

The reconstructed range suitability maps over time were used as an input for CISGeM. In this framework, the genetics of multiple populations can be modelled within a spatially explicit reconstruction of the world where the suitability of each deme changes through time according to the SDM back-cast suitability scores (Fig. 1.). Using an Approximate Bayesian Computation framework, we fitted basic demographic parameters such as population growth rate and migration, as well as the link between SDM suitability scores and local population sizes. The mean pairwise genetic differentiation (π) between all possible pairs of populations were used as summary statistics that had to be matched by the model, thus reconstructing in full the genetic population structure observed at present. We performed a Monte-Carlo sweep of the input parameters (Supplementary Table 1.), generating a total of 60,661 simulations. A power analysis confirmed that we had good power to infer allometric scaling exponent, allometric scaling factor and undirected expansion coefficient, but only limited power for directed expansion coefficient and intrinsic growth rate (Supplementary Fig. 3. & 4., Supplementary Table 2.).

We first confirmed that the model was able to capture the observed genetic patterns. For all of the 210 π between pairs of populations, the empirical value was included in the range produced by the simulations (Supplementary Table 3.). Furthermore, we used Principal Component Analysis to summarise these statistics (see Materials and Methods for details), generating a graphical representation of the empirical data in the context of the simulations: in Fig. 2 D we can see that the model was able to capture the patterns of genetic variation observed in the data in the first two components. We further confirmed that this was the case in all pairwise combinations of the first 4 principal components, which capture ∼ 70% of variance in pairwise π (see Supplementary Fig. 5., and a Scree plot as Supplementary Fig. 6.), ensuring that the model was able to recreate all major aspects of the data simultaneously. Besides checking each pair of components, we also formally tested that the observed data fitted within the 4 dimensional hypervolume formed by the 4 Principal Components (i.e. confirming that all components could be captured simultaneously, thus ensuring that there are no trade-offs in the fit that were missed in the pairwise plots). Finally, we tested model fit with ‘*gfit*’ from the ‘*abc*’ package (30). This test verifies that the distance between the observed and the simulated data is not significantly larger than the distance of a random simulation to other simulations (and thus that the model is able to capture the patterns seen in the data): our model recovered a p value of 0.379 which implies a good fit (Supplementary Fig. 7.). Given these results, we are confident that our model, which is constrained by the range dynamics inferred from the SDM, is able to capture the processes that generated the observed patterns of population variation in yellow warblers.

We used a random forest algorithm (ABC-RF) (31) to generate posterior probabilities of the input demographic parameters given the observed levels of pairwise population differentiation (Supplementary Fig. 8.). The metapopulation dynamics were characterised by limited migration among colonised demes (low values in the undirected expansion coefficient rates *m_r_*, Supplementary Fig. 8. E) in accordance with observations that this species tends to be philopatric in its breeding range. Whilst we were not able to accurately quantify the size of colonisation bottlenecks (defined by the directed expansion coefficient, *m_d_*, which defines the proportion of individuals that move into an unoccupied area, Supplementary Fig. 8. C), the posterior distribution dipped towards large values of this parameter, thus favouring moderate to strong bottlenecks. However, irrespective of the actual strength of these founder events, the low subsequent migration would have preserved the pattern of isolation by distance set up by the sequential founder events along the colonisation routes (see the analysis on common ancestor events for a visualisation of the strength of this effect).

From the top 2.5% best fitting simulations (n=1000 runs with the smallest sum of Euclidean distances between observed and predicted statistics), we reconstructed the demography of the species through space and time. The average demographic profile, calculated as a medians of population size across these simulations, shows that the yellow warbler was forced to contract its range at the peak of the Last Glacial Maximum (∼21kya) as the ice sheets grew across the north of the continent (Fig. 3. A&B). At this point, the population existed in a restricted but broadly continuous range in the south of the continent. As the climate ameliorated, northward range expansion became possible. However, the pattern of recolonization was uneven. By 13kya, our model reconstructs an expansion mostly following the corridor that opened between the Laurentide and Cordilleran icesheets on the west of the continent, whilst expansion on the eastern side was limited (Fig. 3. C). The western spread continued at a pace with the melting of the Cordilleran ice sheet (Fig. 3. D), but the eastern expansion lagged behind due to the slower melting of the Laurentide icesheet (Fig. 3. E). The central and eastern part of the continent were fully colonised by 5kya, when the ice sheets were fully melted (Fig. 3. F).

**Figure 3.**
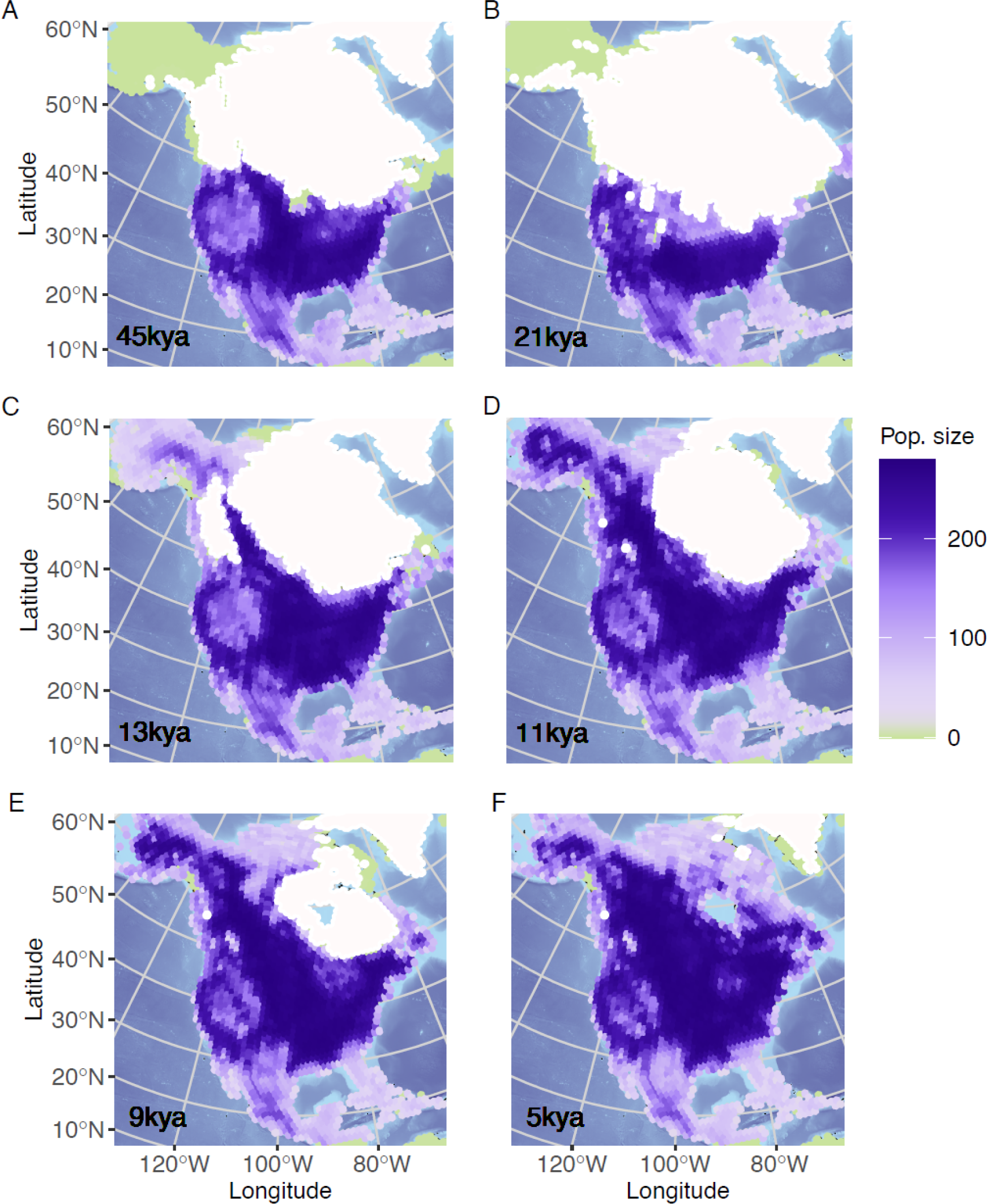
Weighted median population size (per deme) of yellow warbler at A) 45kya, B) 21kya, C) 13kya, D) 11, E) 9kya, F) 5kya, from 1000 simulations retained during the parameter estimation. White regions are areas uninhabitable for yellow warblers at the given point in time.

The importance of this asymmetric expansion in setting up the patterns of genetic diversity across the range of yellow warblers can be seen by mapping geolocation of common ancestor (CA) events that occurred between populations. These events allow us to reconstruct gene flow through time, as shaped by colonisations and subsequent connectivity, revealing how the patterns of diversity have emerged. To track connectivity across the metapopulation, we focus on two populations each from the eastern, central and western part of the continent, and considering two regions at a time, we plotted CA events among their respective populations (Fig. 4. A). When we considered populations from the East and West (Fig. 4. B), we can see that common ancestor events between these two clusters show a “v” shape that matches closely the shape of the ice sheets at 13kya, when the postglacial expansion occurred. Importantly, the same pattern was also found when we considered only West and Central populations (Fig. 4. C), albeit with a greater intensity of events in the corridor between the two populations. Even though we did not have any population from the East, common ancestors event reveal that central populations are linked to that area, thus representing a mix of the western and eastern arms of the expansion. The same is true when we considered only Central and East populations Fig. 4. D). Together with the reconstructed demography in Fig. 3., this pattern shows the importance of the early expansion up the west coast, followed by subsequent expansion up the east coast, in setting up an initial divergence of the clades, which then mixed in the central region comparatively recently. The signature left by the founder events that occurred during the expansion have not yet been eroded by the relatively low levels of migration, explaining the current patterns of genetic diversity and structure in this species.

**Figure 4.**
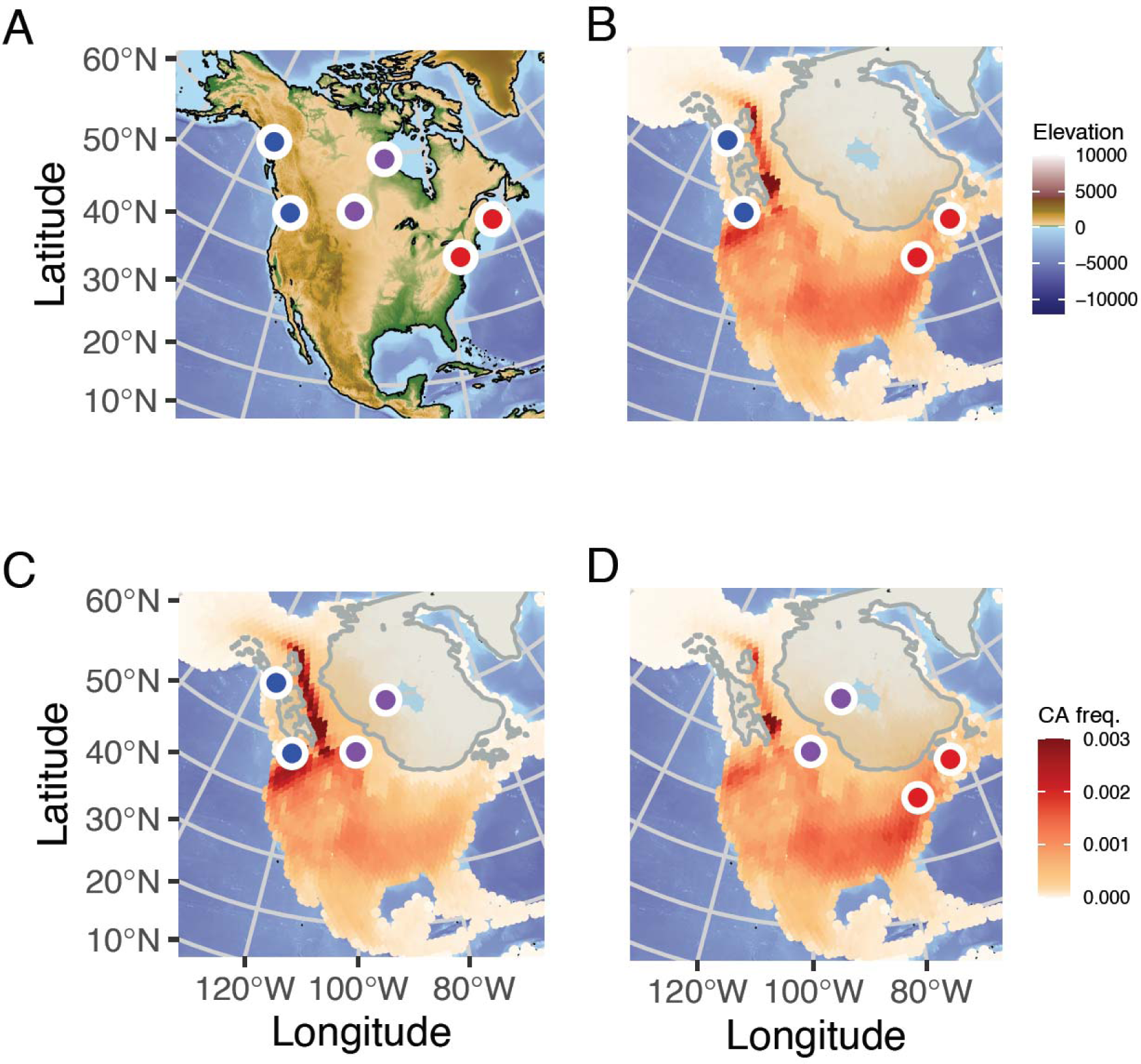
The location of common ancestor (CA) events are plotted across a map of North America. A) is an elevation map of the region with all six sampling locations labelled. In B-D colour density represents proportion of total CA events on the map that occur in each deme. B) is an analysis based on two populations each from West (blue) and East (red) regions, C) is based on West (blue) and Central (purple) regions, and finally D) the Central (purple) and East (red) regions.

## Discussion

In this study, we examined the relative roles of different forces that may have driven modern-day genetic structuring in a widespread species. We used a set of complementary datasets to explore structure in the North American yellow warbler (*Setophaga petechia*), a common passerine species. By integrating genetic data and climatic and environmental variables through time into a spatially-explicit modelling framework (CISGeM), we were able to build a detailed reconstruction of the population dynamics for this species, stretching back through the last fifty thousand years.

Recent work has cautioned against naïve and independent interpretation of either SDMs or genetic based approaches for reconstructing population histories (16), especially when considering species-wide responses. Numerous idiosyncratic factors can confound any single approach but by quantitatively merging genetic data and SDMs our model was able to draw on multiple lines of evidence to reconstruct population size changes, track potential range expansions, and simulate recolonisation dynamics, whilst capturing the genetic structure found in the modern population. With this information, we were able to explore the extent to which expansion dynamics could explain modern genomic patterns of the yellow warbler.

East-west population structure, as found in the yellow warbler, is not an uncommon pattern in North America. These genetic differences, as well as variation in other traits such as migratory behaviour, are often considered to support the existence of isolated refugia during glaciations (e.g. (20,32)). However, recent work on refugia has shown that the patterns of diversity found in the Northern Hemisphere only fit the expectations from cyclic expansion-contraction fragmenting ranges and driving genetic variation at a coarse level (8,9,33).

By explicitly modelling the recolonization dynamics, we have demonstrated a plausible explanation for the formation of genetic structure over time, without the need of multiple glacial refugia. Spatially explicit models have been used to show that range expansions can have profound impacts on the distribution of genetic diversity (12). By reconstructing in details the dynamics of the recolonisation of yellow warblers, we were able to show that, to a large extent, this passerine species tracked the uneven (asynchronous) retreat of the Laurentide Ice Sheet, with a longitudinally unequal progression northward (Fig. 3.). Despite the species exhibiting a single large glacial refugium, the asymmetrical pattern of re-expansion generates the genetic structure of east, west, and central population clusters found in the empirical genetic data. This implies a key role for post-glacial re-expansion in shaping modern-day populations.

The important role of re-expansion dynamics has recently been highlighted in a range of different species e.g. (8,12,33), though it would be naïve to assume that the complex patterns of diversity found in real populations could be easily explained by a single mechanistic process (34). Our work highlights that, at the very least, modern population diversity and structure may have originated from a combination of different processes, each of which needs to be carefully considered.

We acknowledge that, within this framework, we were unable to consider the possible influence of biotic interactions which may have impacted the pattern of recolonization (35). Our model also works with demes that are discrete spatial units of a fixed size, allowing for a step change in the likelihood of common ancestor events occurring within the deme and outside it. Moving away from the discretisation of space could help further ‘naturalise’ our model, and indeed models that incorporate continuous space are rapidly advancing (36). However, there are still major computational challenges to overcome before these tools would be suitable for an area on the scale of this study.

Whilst theories that describe broad patterns have been crucial to increasing our understanding of the likely impacts environmental changes have had on populations, we now realise that North American avifauna is probably a composite of species with different histories (37). Species have responded individually to the rapid climate changes faced in the Pleistocene and therefore we would not wish to claim our findings refute the existence and effect of North American glacial refugia for birds. However, now the resources and techniques exist to study the idiosyncratic responses of different species, and it will be possible to assess the importance of isolated refugia in shaping the genetic structure of species. Furthermore, an increased understanding on the different population dynamics that underlined species responses to the large climatic changes that occurred over the last glacial cycle might provide an important tool to refine our ability to predict the responses of species to anthropogenic change in the future.

## Materials and Methods

### Study species

The North American yellow warbler (*Setophaga petechia*) is a small, riparian, migratory passerine. Today, this common species is widely distributed across the continent. However, despite its large and well-connected contemporary range, the yellow warbler exhibits spatial structure across its range, including multiple mitochondrial clades (20) and clear isolation by distance (18). Although not a species of concern, the yellow warbler has recorded a declining population trend in the North American Breeding Bird Survey between 1966-2015, triggering several studies looking into the species ability to cope in the face of a rapidly changing climate (18,38). One such study by Bay et al. (18) built RAD-seq data from individuals sampled across the species’ range in order to explore potential population trends in response to future climate scenarios. Such data was made available on GenBank and forms the basis of our empirical dataset here.

### Raw genetic data

RAD sequence data for North American yellow warblers (*Setophaga petechia*) from 21 populations (18) were downloaded from the NCBI Sequence Read Archive (SRA). From the 269 accessions associated with the Bay et al. paper we chose to focus on only the individuals included in the original analysis (n = 223), individuals for which full information about their breeding population was available. A further 22 samples were dropped as the file sizes were under 75MB and, therefore, were likely to have low coverage. One final exclusion was made, GenBank accession number SRR6366039, as the sample was found to be an outlier with a measure of diversity higher than the range of all other samples, despite comparable levels of coverage and number of sites. This left 200 samples for further analysis. These individuals were sampled from across the modern population range, providing a good overview of the population genetics of this species, see Fig 2 A. for sampling locations.

### Clustering analysis

RAD-seq methods are known to create specific biases in estimated allele frequencies, potentially affecting downstream analysis of the data (39). Using allele frequencies derived directly from the sequence data in a genotype-free method has been shown to account for RAD-seq specific issues, improving population genetic inferences (39). Therefore, we used Analyses of Next-Generation Sequencing Data (ANGSD) (40,41) to infer genotype likelihoods directly from aligned BAM files. Filters were set to only include SNPs with a p value of < 2 x10^−6^ and only keep sites with at least 100 informative individuals. These ANGSD genotype likelihood values were then used as input for NGSadmix to calculate population admixture, setting a *K* (presumed cluster number) value of 2 and keeping minimum informative individuals at 100.

### Observed genetics for CISGeM

In order to calculate pairwise π (the average number of differences between two sequences, normalised by the number of available positions), we first calculated genotype likelihoods in ANGSD. Input files were aligned BAM files, we used the samtools genotype likelihood method and inferred the major and minor allele from these likelihoods, with the command below:

angsd -GL 1 -out genolike -doGlf 1 -doMajorMinor 1 -bam bam.filelist

We then computed pairwise π from the ANGSD output; since our population genetic simulations (see below) modelled haploid samples (as it is the case of most genetic simulators, e.g. msprime (42)), we used the below formula:

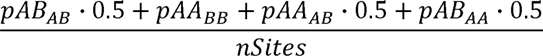

In order to make the modelling computationally feasible, we then investigated how many samples were needed to get a reliable estimate of π for each population (Supplementary Fig. 9.). This analysis showed that five diploid individuals, or ten chromosomes, provided a reasonable compromise for noise. All estimates of pairwise π were therefore re-computed with only five individuals per population. As estimates were consistent with the values from the full dataset (Supplementary Fig. 10.), for computation efficiency of the model, all future analyses were based on this subset of the data.

### Species Distribution Modelling for world reconstruction

The range and population size of a species changes in time and space according to fluctuations in resources and environmental conditions. In order to build a spatially explicit model it is first necessary to use Species Distribution Modelling (SDM) to reconstruct how population ranges and demographics may have changed through time. We used the biomod2 (43) library in R (44) to fit the SDM; the code is available as a Rmarkdown document in the supplementary material (Supplementary Materials Pipeline 1.).

### Climate reconstructions

Climate data for North America were drawn from a 0.5° resolution dataset for 19 bioclimatic variables; Net Primary productivity (NPP), Leaf Area Index (LAI) and all the BioClim variables (45) with the exclusion of BIO2 and BIO3; covering the last 50,000 years in 1,000 year time steps from the present to 22kya and in 2,000years time steps before that date (46). This dataset was originally constructed from a combination of HadCM3 climate simulations of the last 120,000 years (47), high-resolution HadAM3H simulations of the last 21,000 years (48), and empirical present-day data. The data had been downscaled and bias-corrected using the Delta Method (49). Bioclimatic variables through time were then used as input data to inform the SDM.

### SDM data preparation

Species occurrences data for the present day were initially downloaded from the GBIF database (https://www.gbif.org), the original downloads are available at the following DOI: S. petechia 10.15468/dl.jfkwcg (GBIF.org). These data were then filtered based on the attributed accuracy of the coordinates (maximum error: 1 km) and additionally, only points that were within Birdlife breeding and resident geographical ranges (22) were retained. Remaining occurrences were then matched to the 0.5° resolution grid used for the palaeoclimatic reconstructions and, as the method works on presence/absence data and not frequency, only one presence per grid cell was kept.

This cleaned observation dataset was then used to define a set of informative bioclimatic variables with the most influence on the species distribution for use in the Species Distribution Model (SDM), through visual check of how much the distribution of the variable values differed between the observation points and the whole area. We selected the variables with highest differences between the two curves, which are most likely to be relevant for the species, and then, in order to avoid using highly correlated variables, which may increase noise in the data, we constructed a correlation matrix between the chosen variables associated with each of the retained observations. Where two values were highly correlated, the variable with the lowest overall correlation across the matrix was kept, allowing us to select a set of uncorrelated variables (threshold = 0.7) leaving us with the following ones to be used for SDM modelling: LAI (leaf area index), BIO7 (Temperature Annual Range), BIO8 (Mean Temperature of Wettest Quarter), BIO14 (Precipitation of Driest Month).

Geographic biases in sampling effort are common when observation data are collected opportunistically, which is the case for the GBIF database. In order to reduce this bias, we thinned our dataset using the R package *spThin* (50) enforcing a minimum distance of 70 km between observations. Given the random nature of removing nearest-neighbour data points, we repeated this step 100 times (‘rep’ = 100) retaining for further analysis the result with the maximum number of observations after thinning.

### SDM modelling

The SDM was built with the R package biomod2 (43) following the same procedure used in Miller et al. (51). The thinned observation dataset was used as presences whilst the landmass of North America was considered as background. The same number of pseudo-absences as presences were then drawn five separate times, at random, from outside the BirdLife resident and breeding masks: creating five independent datasets for analysis. For each data set, following Bagchi et al. (52), models were then run independently using four different algorithms: generalised linear models (GLM), generalized boosting method (GBM), generalised additive models (GAM), and random forest.

Spatial cross-validation was used to evaluate the model; 80% of the data were used to train the algorithm and the remaining 20% to test it. Initially, both the presences and the five pseudoabsences datasets were subdivided in 14 latitudinal bands using the R package BlockCV (53). Each band was given a ‘band ID number’, looping sequentially through numbers 1-5 until all bands were labelled. Then the bands were assembled into five working data splits grouped by their band ID (numbers 1-5). This was performed to maximise the probability of having at least some presences in all five data splits as a data split cannot be used for evaluation if it contains only absences. Each of the four models (GLM, GBM, GAM, and random forest) were then run five times (once for each pseudoabsence run), using in turn four of the five defined data splits to calibrate and one to evaluate based on TSS (threshold = 0.7).

Finally, a full ensemble combining all algorithms and pseudoabsences runs (54) was created, using only models with TSS > 0.7, averaged using four different statistics: mean, median, committee average and weighted mean. The statistic showing the highest TSS, the mean, was then used to predict the probability of occurrence in each grid cell. This was then projected for all available time slices from the present to 50 thousand years ago.

### CISGeM Demography

CISGeM’s demographic module consists of a spatial model that simulates long-term and global growth and migration dynamics of yellow warblers. These processes depend on a number of parameters (see Supplementary Table 1.), which we later estimate statistically based on empirical genetic data.

The model operates on a global hexagonal grid of 40962 cells that represent the whole world (the distance between the centre of two hexagonal cells is 241 ±15 km); 2422 grid cells make up North America. Each time step represents 1 year, the generation time of yellow warblers. Each time step of a simulation begins with the computation of the carrying capacity of each grid cell, i.e. the maximum number of individuals theoretically able to live in the cell on the basis of the environmental resources at the given point in time. Here, we estimate the carrying capacity in a grid cell *x* at a time *t* as

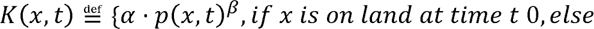

where *p*(*x,t*) denotes the probability of a species inhabiting cell *x* at time *t* (see section ‘Species Distribution Modelling’). The particular function used here was chosen based on analysis of SDM projections and census data of Holarctic birds (R. Green, pers. comm.).

The estimated carrying capacities are used to simulate spatial population dynamics as follows. We begin a simulation by initialising a metapopulation of yellow warblers covering the largest set of adjacent inhabitable demes within their potential distribution 50k years ago (defined as *t_0_*), with each deme at carrying capacity (given the climate 50k years ago and the set of parameters used in a given simulation).

At each subsequent time step between *t*_0_ and the present, CISGeM simulates two processes: the local growth of populations within grid cells, and the spatial migration of individuals across cells. We used the logistic function to model local population growth, estimating the net number of individuals by which the population of size *N*(*x,t*) in the deme *x* at time *t* increases within the time step as

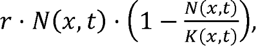

where *r* denotes the intrinsic growth rate. Thus, growth is approximately exponential at low population sizes, before decelerating, and eventually levelling off at the local carrying capacity.

Across-cell migration is modelled as two separate processes. First, we have a non-directed, spatially uniform movement into all neighbouring grid cells, which represents the level of migration that we would expect in a fully saturated environment (i.e. where all grid cells are at carrying capacity). The second is a directed movement along a resource availability gradient on the other hand; note that this type of preferential movement does not require complete knowledge of the environment, it can be observed even if animals explore randomly but stop preferentially in areas with more available resources (i.e. where the local population is below carrying capacity). Under the first movement type, the number of individuals migrating from a cell *x* into any one of the up to six neighbouring cells is estimated as

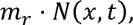

where *m_r_* is a mobility parameter. This mechanism is equivalent to a spatially uniform diffusion process, which has previously been used to model random movement in other species (55). Under the second movement type, an additional number of individuals moving from a grid cell *x*_1_ to a neighbouring cell *x*_2_ is estimated as

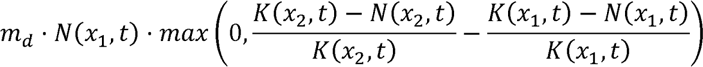

where *m_d_* is the directed mobility parameter. The number 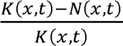 represents the relative availability of unused resources in the cell *x* at time *t*, equalling 1 if all natural resources in *x* are potentially available for yellow warblers (*N*(*x,t*) = 0), and 0 if all resources are used (*K*(*x,t*)). Thus, individuals migrate in the direction of increasing relative resource availability, and the number of migrants is proportional to the steepness of the gradient. The distinction between directed and non-directed movement allows us to capture the higher fraction of migrants that move into empty or lightly inhabited demes (in some other models, e.g.(15), this is terms the colonisation rate when representing the first time step of colonisation, but in our representation we allow this increased mobility to persist as long as a grid cell is below carrying capacity).

For some values of the mobility parameters *m_r_* and *m_d_*, it is possible for the calculated number of migrants from a cell to exceed the number of individuals in that cell. In this scenario, the number of migrants into neighbouring cells are rescaled proportionally such that the total number of migrants from the cell is equal to the number of individuals present.

Similarly, it is in principle possible that the number of individuals present in a cell after all migrations are accounted for (i.e., the sum of local non-migrating individuals, minus outgoing migrants, plus incoming migrants from neighbouring cells) exceeds the local carrying capacity. In this case, incoming migrants are rescaled proportionally so that the final number of individuals in the cell is equal to the local carrying capacity. In other words, some incoming migrants perish before establishing themselves in the destination cell, and these unsuccessful migrants are not included in the model’s output of migration fluxes between grid cells. In contrast, non-migrating local residents remain unaffected in this step. They are assumed to benefit from a residential advantage (56), and capable of outcompeting incoming migrants.

CISGeM’s demographic module outputs the number of individuals in each grid cell, and the number of migrants between neighbouring grid cells, across all time steps of a simulation. These quantities are the used to reconstruct genetic lineages.

### CISGeM predicted genetics

Once a global population demography has been constructed, gene trees are simulated. This process is dependent on the population dynamics recorded in the demography stage and assumes local random mating according to the Wright-Fisher dynamic. At present, individuals are randomly sampled to represent the available genomes, and their ancestral lines are tracked back through the generations, recording which cell each line belongs to. Every generation, the lines are randomly assigned to a gamete from the individuals within its present cell. If the assigned individual is a migrant or coloniser, the line moves to the cell of origin for that individual before ‘reproduction’. Whenever two lines are assigned to the same parental gamete, this is recorded as a coalescent event, and the two lines merge into a single line representing their common ancestor. This process is repeated until all the lineages have met, reaching the common ancestor of the whole sample. If multiple lineages are still present when the model reaches the generation at which the demography was initialised, the lines enter a single panmictic ancestral population of size *K*_0_ until sufficient additional coalescent events have occurred for the gene tree to close (thus transitioning from a discrete time W-F model to a coalescent model).

### Parameter exploration

Parameter space was explored with a Monte Carlo sweep in which demographic parameters were randomly sampled from flat prior ranges: directed expansion coefficient [0.0,0.14], undirected expansion coefficient [0.0,0.04], intrinsic growth rate [0.02,0.15], allometric scaling exponent [0,1.5], and allometric scaling factor [20,5000] on a log_10_ scale. A fixed mutation rate of 2.3×10^−9^ μ/Site/Year was used (57). We run 60,661 simulations, of these 43,387 had a demography such that all sampling locations were reached, and thus produced genetics summary statistics.

### Confirming that the model fits the data

We confirmed that the model was able to realistically capture the observed genetics by first testing that the observed values of all 210 summary statistics fell within the range generated by the model. When fitting models by linear ABC, it is customary to check all pairwise combinations of summary statistics to ensure that there are no tradoffs that prevent fitting certain quantities simultaneously. When using large numbers of summary statistics as done for random forest ABC, this is not practical (and arguably not necessary). However, to make sure that our model could capture all key aspects of the genetics patterns simultaneously, we used a similar approach using the first 4 principal components (PC) of the summary statistics (which explains 70% of the total variance and were shown by a Scree plot to be a suitable number as further components explained little additional variance; see Supplementary Fig. 5. and Supplementary Fig. 6.). Besides a visual check, we also tested formally (by setting up an appropriate simplex) that the observed values in the first PC axes fell within the 4-dimensional hyperspace formed by all the simulations (i.e. that all 4 PCs could be captured simultaneously).

Finally, we further confirmed the quality of the model fit using a formal hypothesis testing approach based on randomisation from the R package ‘*abc*’ (30). The ‘*gfit’* function (58) estimates a goodness of fit test statistic, or D-statistic, as the median Euclidean distance between the observed summary statistics and the nearest simulated summary statistics. For comparison, a null distribution of D is then generated from summary statistics of 1000 pseudo-observed datasets (i.e. a random set of 1000 simulations). A goodness of fit p-value can then be calculated as the proportion of D based on pseudo-observed data sets that are larger than the empirical value of D. Consequently, a non-significant p-value signifies that the distance between the observed and accepted summary statistics is not larger than the expectation, confirming that the model fits the observed data well, (Supplementary Fig. 7.).

### ABC parameter estimation

Parameter estimation was performed with an ABC random forest (RF) approach implemented via the R package ‘*abcrf’* (31,59). Forests of 1,000 trees were used, fitted to all possible 210 pairwise pi among populations. We performed a power analysis by taking out 1000 simulations as a test set, and then using the remaining simulations to train the forests for each demographic parameter. We assessed the power of the analysis by looking at both the predicted mean and median value for each of the parameters in the 1000 sets, and estimated the R^2^, the median of the standardised error and of the square root of standardised error, and the coverage. Results based on the median and mean of the posterior were equivalent (Supplementary Fig. 3. and Supplementary Fig. 4.): we had good power for allometric scaling factor, allometric scaling exponent and undirected expansion coefficient, whilst we had little ability to infer directed expansion coefficient and intrinsic growth rate (Supplementary Table 2.).

## Supporting information

Species Distribution Modelling pipeline

## Author contribution

E.F.M. and A.M. devised the project. E.F.M. and Z.X. ran the genetic analysis under the supervision of P.M.D. and A.M.. M.L. ran the Species Distribution Modelling. M.K., R.B., and A.M. developed the modelling software with help from E.F.M, P.M.D. and G.L.S.. M.S. helped with the interpretation of results. E.F.M and A.M. wrote the manuscript with feedback from all the co-authors.

