## Supplementary material for "Post-glacial expansion dynamics, not refugial isolation, shaped the genetic structure of a migratory bird, the yellow warbler"

Supplementary Materials

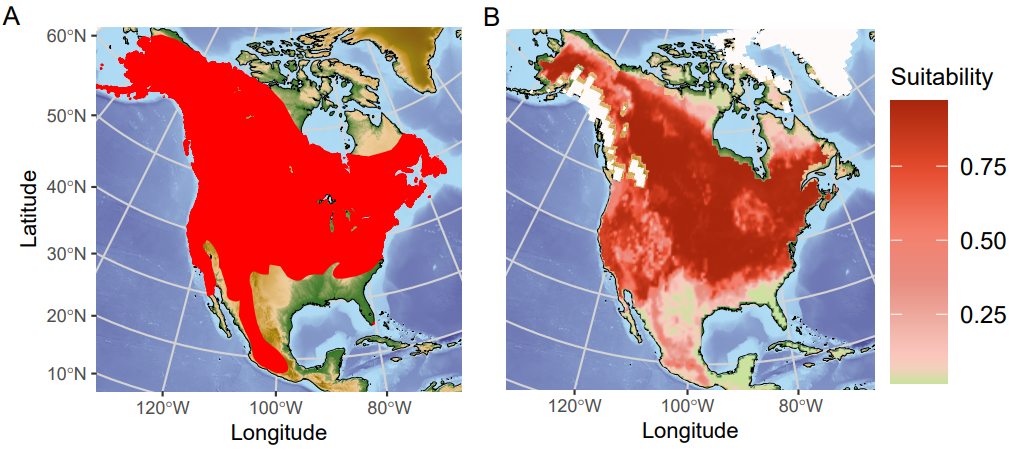

**Supplementary Figure 1**. A) shows the best estimated breeding and resident geographical range of the North American yellow warbler at the present day according to BirdLife. B) shows the potential species range for the same time point, reconstructed with our SDM. As we are only considering climatic suitability in our SDM we do not explicitly exclude bodies of fresh water such as the Great Lakes from the estimates, unlike the BirdLife range. Nonetheless, the predicted potential distribution matches well the best range estimates for the species.

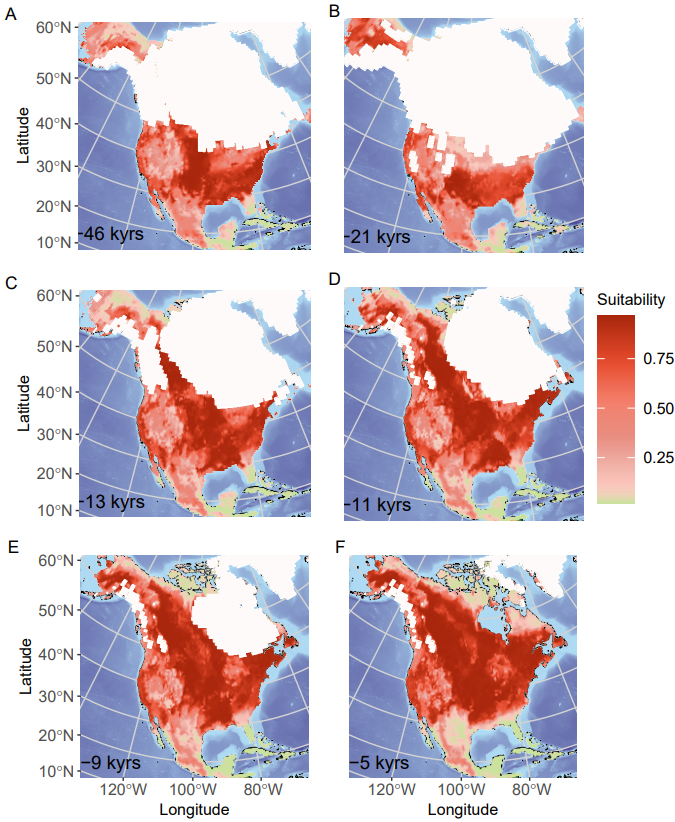

**Supplementary Figure 2.** Plots showing potential range for the North American Yellow Warbler through time.

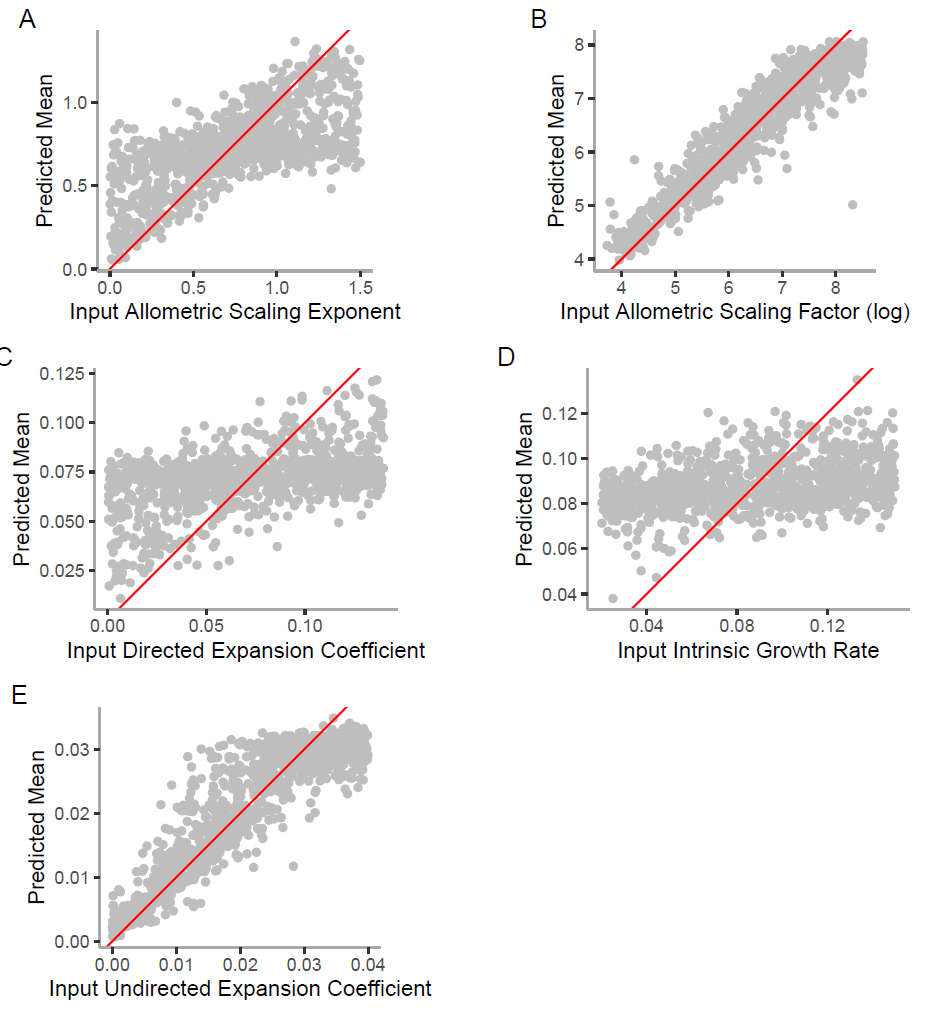

**Supplementary Figure 3.** Input vs predicted mean values for the demographic parameters as obtained from the power analysis (n=1000). A good power is indicated by a 1 to 1 relationship.

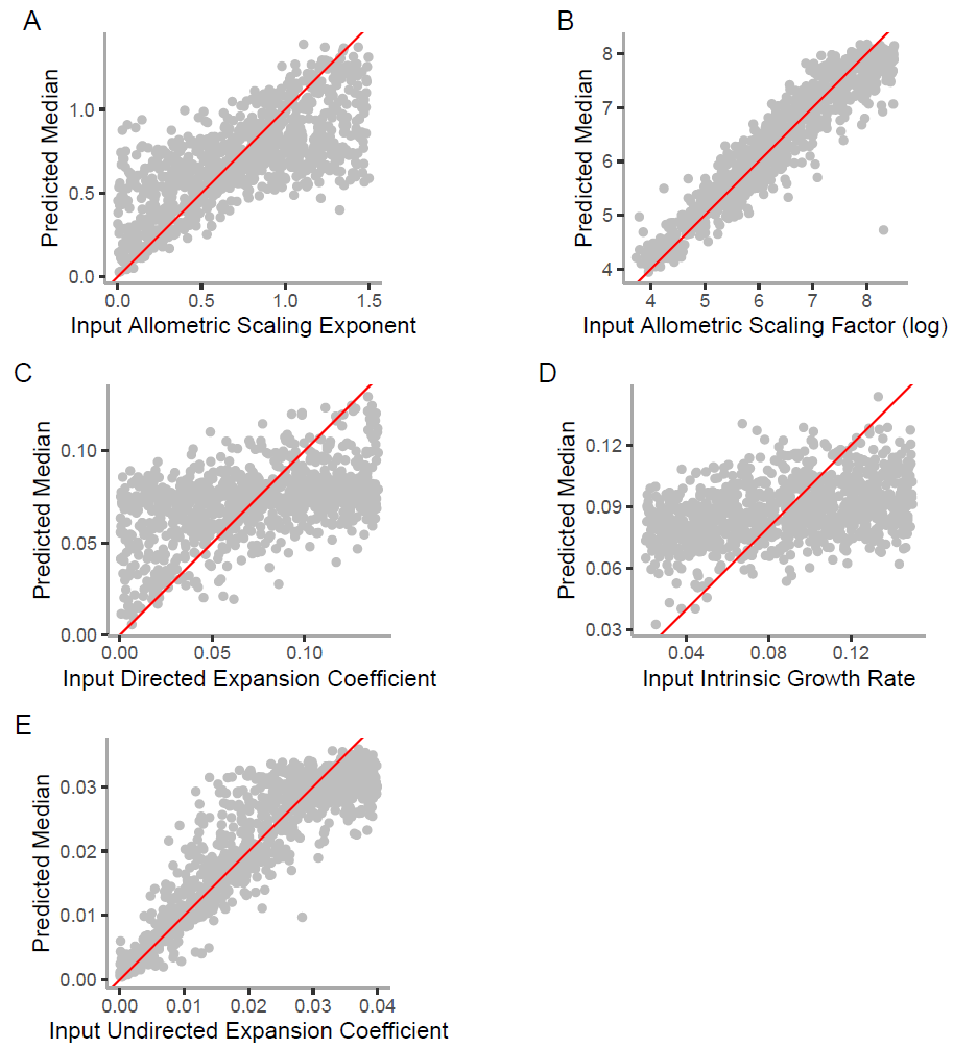

**Supplementary Figure 4.** Input vs predicted median values for the demographic parameters as obtained from the power analysis (n=1000). A good power is indicated by a 1 to 1 relationship.

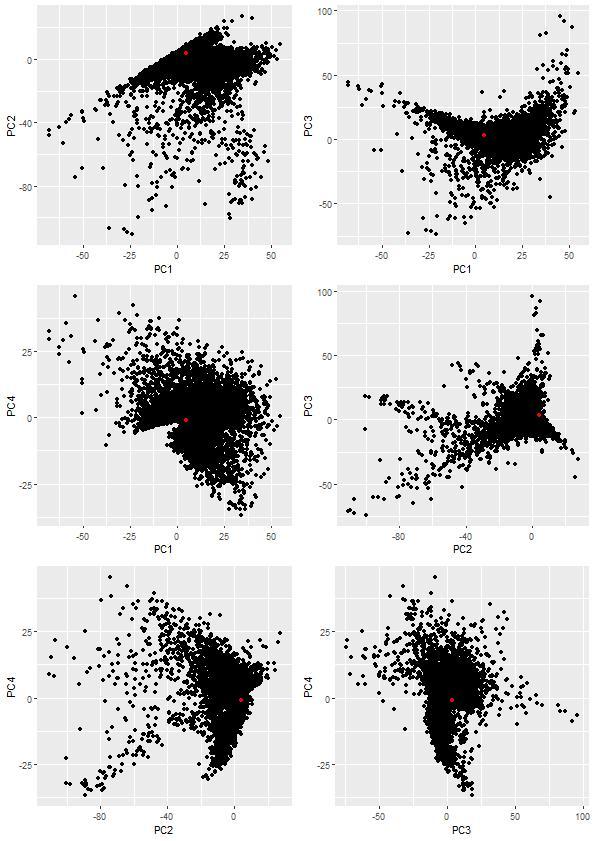
**Supplementary Figure 5.** Pairwise plots of the first 4 PCs of summary statistics distributions from a Monte-Carlo sweep. Observed values are in red, simulated values are in black.

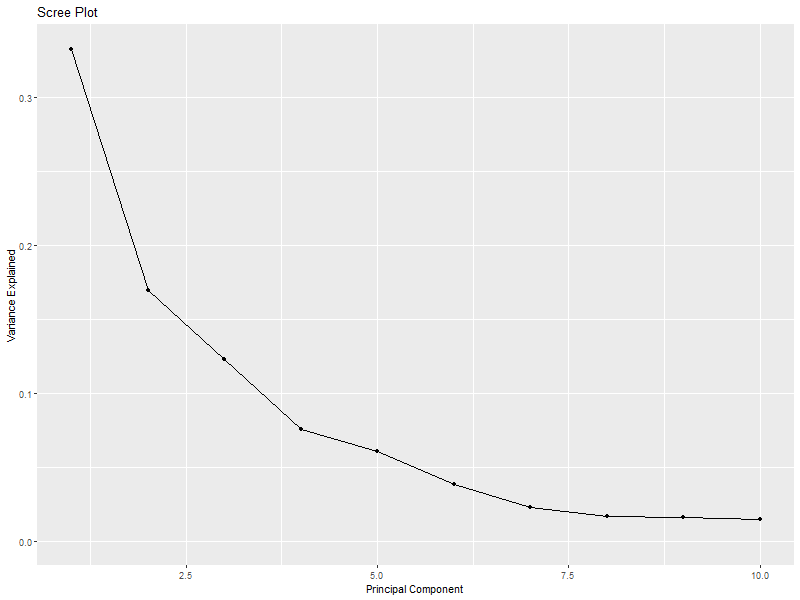

**Supplementary Figure 6.** The scree plot shows the top 10 PCs and the variance they explained. Top 4 PCs explained 70.0% of total variance.

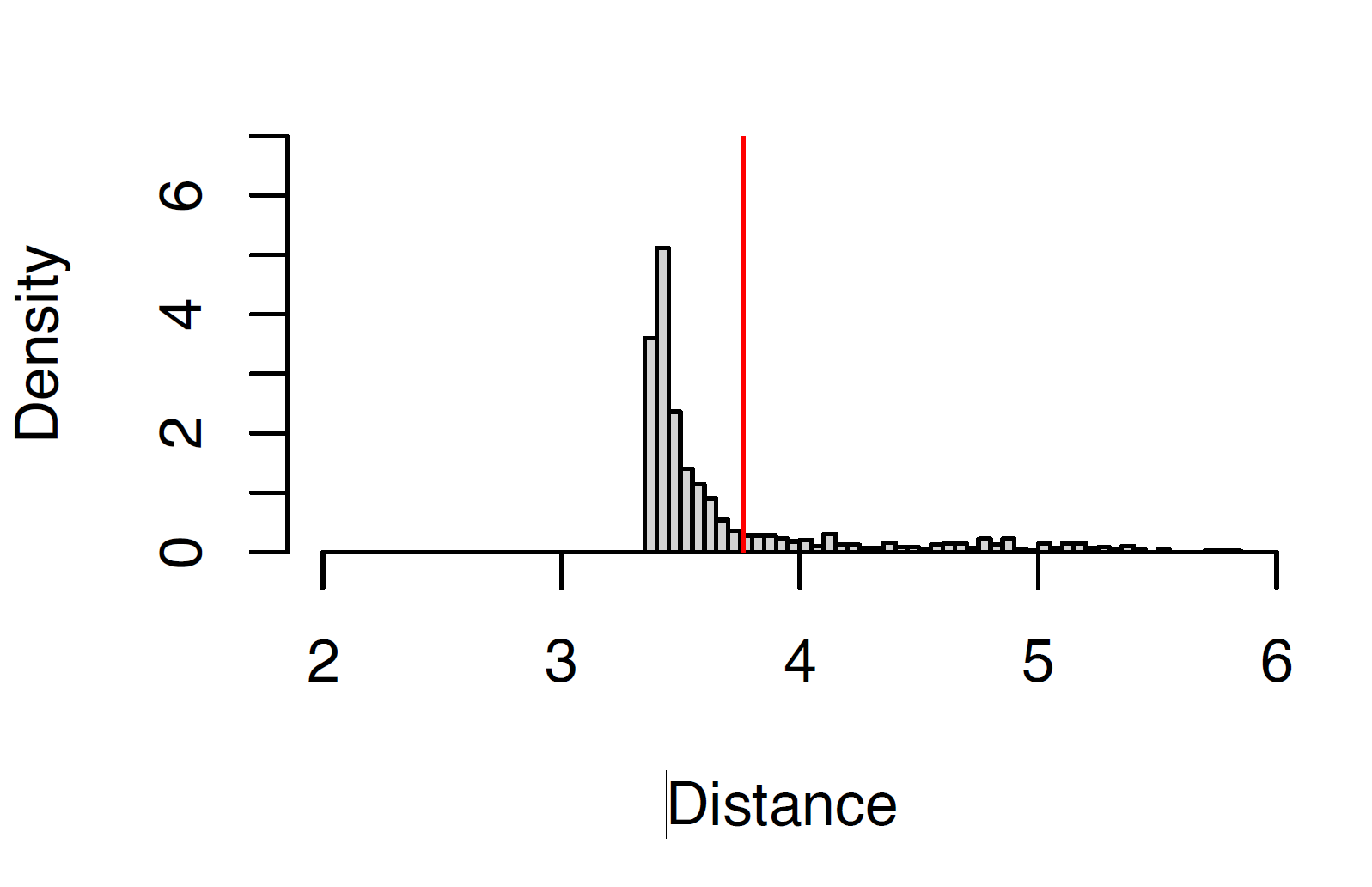

**Supplementary Figure 7.** Histogram of the null distribution of the test statistic for goodness of ﬁt. The red line indicates the observed distance, which is well within the expected range, thus indicating a good fit of the model to the data.

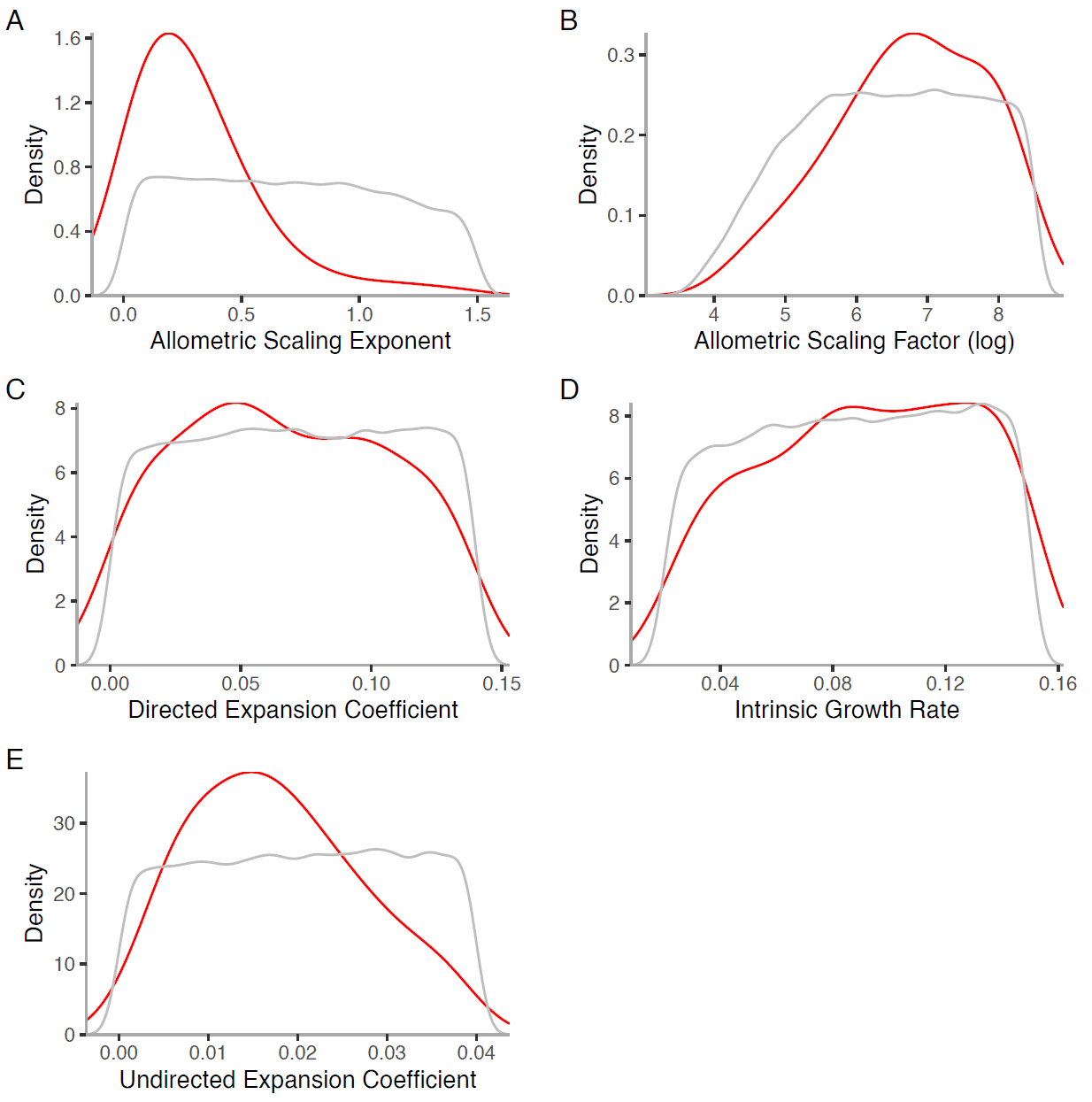

**Supplementary Figure 8.** Posterior distribution of key parameters. Briefly; A and B, the allometric scaling exponent and the allometric scaling factor, determine the shape of the function linking suitability from the SDM to effective population sizes (see Materials and Methods for details). Directed expansion (C), is when individuals migrate in the direction on increasing relative resource availability. Intrinsic growth rate (D) is the local growth rate modelled by a logistic function. Then E is undirected expansion, a spatially uniform movement into neighbouring cells. Grey line shows prior values, solid red line represents the posterior values from abc random forest.

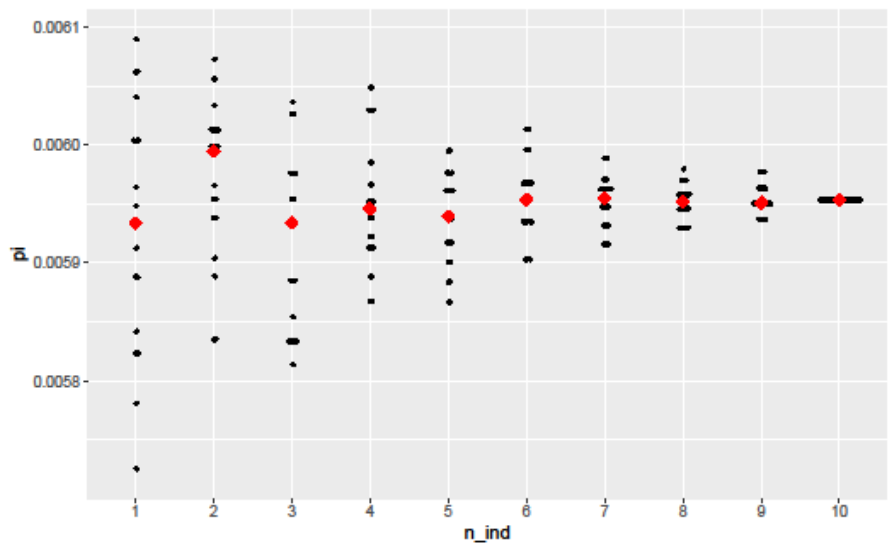

**Supplementary Figure 9.** Plot of pairwise π estimates between two populations. Black dots represent the 20 replicates for each number of individuals, points in red show the median value.

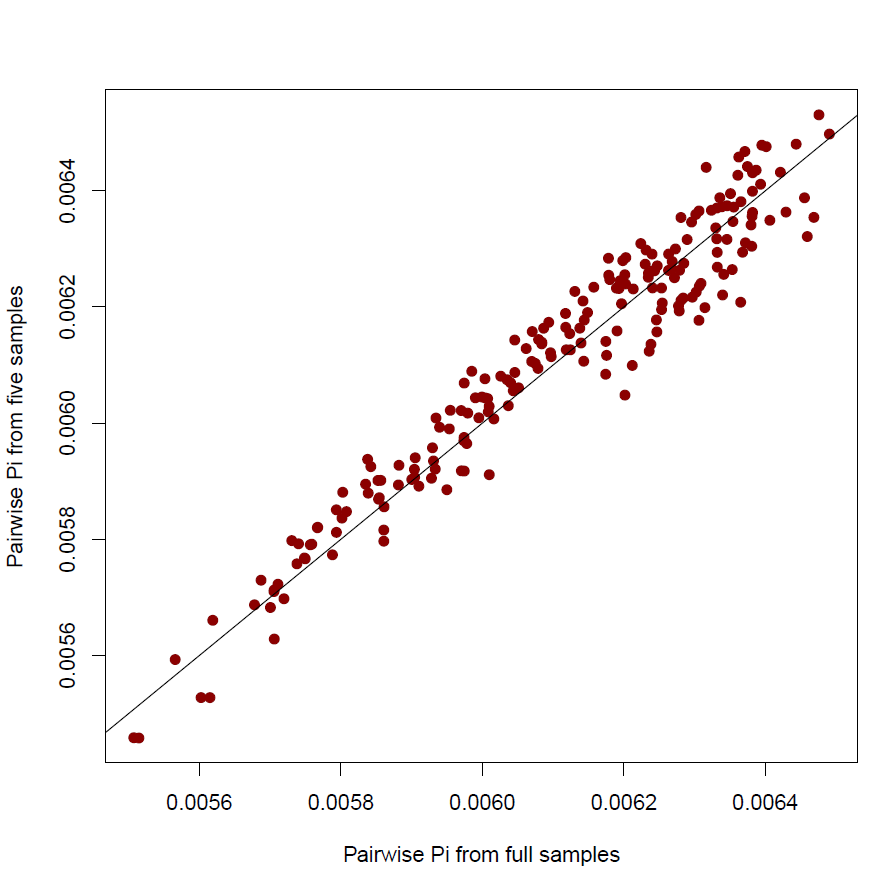

**Supplementary Figure 10.** Relationship between pairwise π calculated from full sample sizes and pairwise π calculated from five individuals per population**.**

| $\alpha$ | Allometric scaling factor for population size | 20 – 5000 |
| --- | --- | --- |
| $\beta$ | Allometric scaling exponent for population size | 0 – 1.5 |
| $r$ | Intrinsic growth rate | 0.02 – 0.15 |
| $m_{r}$ | Non-directed (random) mobility parameter | 0.0 – 0.04 |
| $m_{d}$ | Directed mobility parameter | 0.0 – 0.14 |

**Supplementary Table 1.** Details of parameters used in CISGeM.

| Param | Median of std error of mean posterior | Median of std error of median posterior | Root of Median of Square std error of mean posterior | Root of Median of Square std error of mean posterior | R^2^ of mean | R^2^ of median | cov95 |
| --- | --- | --- | --- | --- | --- | --- | --- |
| Allometric Scaling Exponent | 0.019 | -0.008 | 0.242 | 0.231 | 0.50 | 0.52 | 0.971 |
| Allometric Scaling Factor (Logged) | 0.006 | 0.003 | 0.041 | 0.038 | 0.89 | 0.89 | 0.989 |
| Directed Expansion Coefficient | 0.012 | 0.007 | 0.365 | 0.344 | 0.23 | 0.24 | 0.958 |
| Intrinsic Growth Rate | -0.001 | 0.004 | 0.297 | 0.297 | 0.13 | 0.13 | 0.952 |
| Undirected Expansion Coefficient | 0.006 | -0.008 | 0.157 | 0.150 | 0.80 | 0.81 | 0.977 |

**Supplementary Table. 2.** The table shows the R^2^, the median standardised error and square root of the median standardised error, and 95% coverage for each fitted parameter, based on both the mean and median of the posterior from the ABC-RF.

**Supplementary Table. 3. (In separate xlsx table)** The table shows the observed summary statistics value, min, max and quantiles of simulated summary statistics values, and whether observed value falls into the range of simulated values.
