## Supplementary material for "Post-glacial expansion dynamics, not refugial isolation, shaped the genetic structure of a migratory bird, the yellow warbler": Species Distribution Modelling pipeline

### SDM analyses of Setophaga petechia

Michela Leonardi, Department of Zoology, University of Cambridge.

2021-05-10

#### Introduction

The following code has been used to analyse GBIF data from the American Yellow Warbler *Setophaga petechia* (former *Dendroica petechia*). The first chapter describes the code used to clean and format the species data from the *.csv format downloaded from the GBIF database for the analyses performed. The second chapter details a set of ecological analyses needed to better understand the effect of different climatic variables on the distribution of the species. Those latter are also preparatory for the species distribution modelling (third chapter) performed with the package biomod2.

#### Load and prepare the species data

Setting the working directory and variables, load a low resolution world map

```
library(rworldmap)
# Load world map
newmap <- getMap(resolution="low")

ID <- "S.petechia"

# color palettes
IDcolor <- "darkgoldenrod4"
colBase <- "lemonchiffon1"
cS <- colorRampPalette(c(colBase,IDcolor))(13)
rasterColors <- colorRampPalette(c(colBase,IDcolor))(4)
col <- cS[8]
gray <- "gray58"

# folders
# climate input files
climdir <- "C:/Users/miche/Documents/Progetti/Utilities/Climate/Climate_20200430/NAmerica/"
# ice masks
icedir <- "C:/Users/miche/Documents/Progetti/Utilities/Climate/Climate_20200430/old_Ice_masks/"

# vector of variable names
vars <-c("npp","lai","BIO1","BIO4","BIO5","BIO6","BIO7","BIO8","BIO9",
         "BIO10","BIO11","BIO12","BIO13","BIO14","BIO15","BIO16","BIO17","BIO18","BIO19")
```

A database with recorded presences of the species has been downloaded from GBIF (GBIF.org (19th November 2018) GBIF Occurrence Download https://doi.org/10.15468/dl.jfkwcg) and is available at this link.

Reading the file will give an error message that can be ignored.

```
db <- data.table::fread("0005643-181108115102211.csv") #GBIF file
```

The database contains a total of 1573147 observations. Some of them have one or more "issues" reported in the corresponding column. The following code compares the number and the distribution of observations without issues and with so-called "rounded coordinates".

```
# to check which kind of issues the data has
issue <- unique(db$issue)
#issue[1:4]

# remove all lines with reported issues and above the equator
db1 <- db[db$issue=="" & db$decimalLatitude > 0,]
# remove lines where uncertainty of coordintes is > 1000
db1b <- db1[db1$coordinateUncertaintyInMeters < 1000 |
              is.na(db1$coordinateUncertaintyInMeters),]

# keep lines with "coordinates rounded"
db2 <- db[db$issue%in%issue[c(1,2)] & db$decimalLatitude > 0,]
# remove lines where uncertainty of coordintes is > 1000
db2b <- db2[db2$coordinateUncertaintyInMeters < 1000 |
              is.na(db2$coordinateUncertaintyInMeters),]
```

Plot of a map comparing observations with and without rounded coordinates; they contain respectively 1526468 and 253745 observations. The extent of the region of interest (North America) is between -180°E, -15°E, 8°N, 90°N.

```
# plot map with and without rounded coordinates
png(paste(ID, "GBIF_maps.png", sep="_"), height=6, width=10,  units='in', res=600)
par(mfrow=c(1, 2),      # 1x2 layout
    oma=c(0, 0, 5, 0), # rows of text at the outer bottom left top right margin
    mar=c(3, 1, 3, 1), # space for row of text at ticks and to separate plots
    mgp=c(2, 1, 0))     # axis label at 2 rows distance, tick labels at 1 row

plot(newmap, col="cornsilk", bg="lightblue1", lwd=0.05, border="grey",
     xlim=c(-180,-15), ylim=c(8,90),
     main=paste("Clean data\n", dim(db1b)[1], "observations", sep=" "))#,
```

```
## Warning in wkt(obj): CRS object has no comment
```

```
points(as.numeric(db1b$decimalLongitude), as.numeric(db1b$decimalLatitude), pch=".")

plot(newmap, col="cornsilk", bg="lightblue1", lwd=0.05, border="grey",
     xlim=c(-180,-15), ylim=c(8,90),
     main=paste("Data with rounded coord\n", dim(db2b)[1], "observations", sep=" "))#,
```

```
## Warning in wkt(obj): CRS object has no comment
```

```
points(as.numeric(db2b$decimalLongitude), as.numeric(db2b$decimalLatitude), pch=".")

title(main=paste("GBIF", ID, "database"),cex.main= 3,
      outer=TRUE, line=2)

dev.off()
```

For the analyses only the summer (native breeding) range has been kept. From the Handbook of the Birds of the World website: "Eastern populations leave breeding grounds early, from mid-July, and move South on broad front through North America and return migration also early, reaching breeding grounds from early April in South, late May in far North. Western populations migrate a few weeks later, in both autumn and spring."

A first filter has been applied based on the month of the observation, considering the narrower temporal

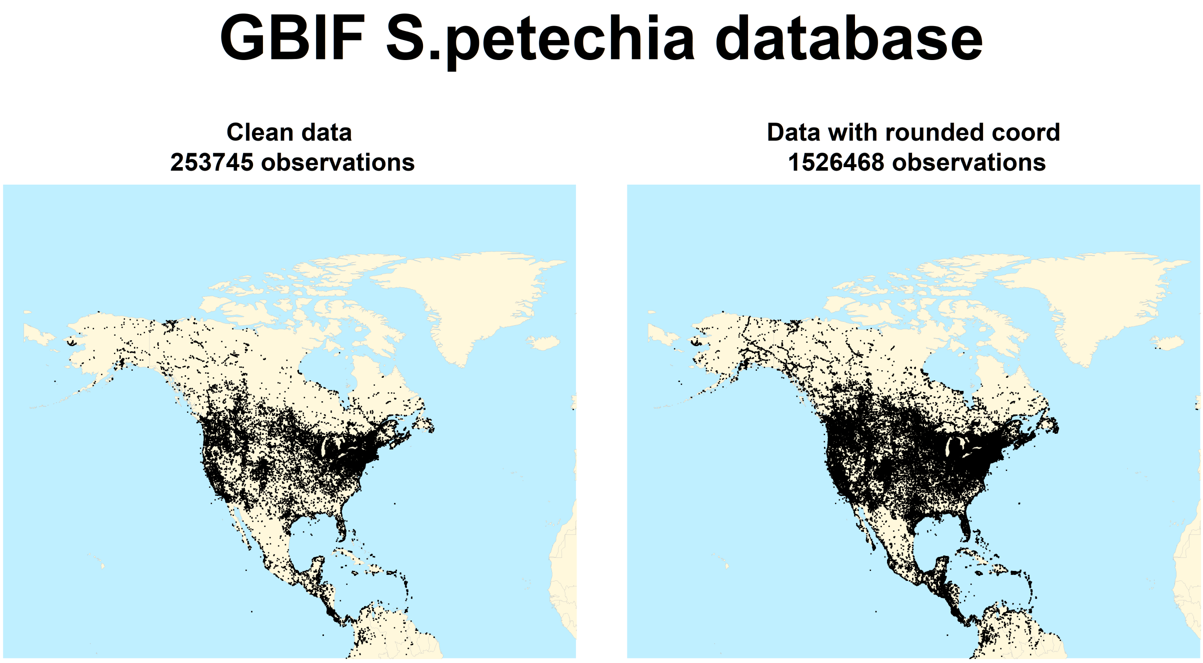


Figure 1: Data without issues vs the same data including also observations with rounded coordinates

range described above

```
months <- c(5,6,7)
sum2 <- db1b[db1b$month %in% months,]

png(paste(ID, "allVSsummer_maps_2.png", sep="_"), height=6, width=10,  units='in', res=600)
par(mfrow=c(1, 2),      # 1x2 layout
    oma=c(0, 0, 5, 0), # rows of text at the outer bottom left top right margin
    mar=c(3, 1, 3, 1), # space for row of text at ticks and to separate plots
    mgp=c(2, 1, 0))    # axis label at 2 rows distance, tick labels at 1 row

plot(newmap, col="cornsilk", bg="lightblue1", lwd=0.05, border="grey",
     xlim=c(-180,-15), ylim=c(8,90),
     main=paste("Clean data\n", dim(db1b)[1], "observations", sep=" "))
```

```
## Warning in wkt(obj): CRS object has no comment
```

```
points(as.numeric(db1b$decimalLongitude), as.numeric(db1b$decimalLatitude), pch=".")

plot(newmap, col="cornsilk", bg="lightblue1", lwd=0.05, border="grey",
     xlim=c(-180,-15), ylim=c(8,90),
     main=paste("Summer data\n", dim(sum2)[1], "observations", sep=" "))
```

```
## Warning in wkt(obj): CRS object has no comment
```

```
points(as.numeric(sum2$decimalLongitude), as.numeric(sum2$decimalLatitude), pch=".")

title(main=paste(ID, "summer observations"),cex.main= 3,
      outer=TRUE, line=2)
```

```
dev.off()
```

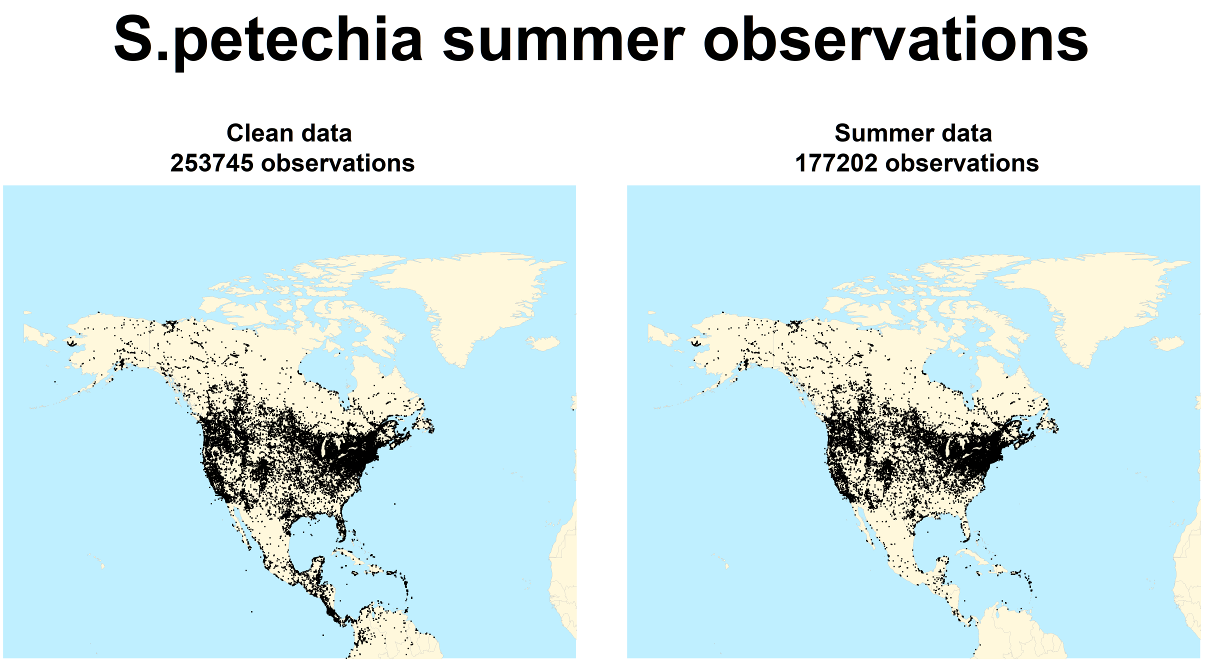


Figure 2: All data VS data only collected during summer

Such filtering reduced the database to 177202 observations. The plot still shows observations outside the expected native breeding range for the species, as reported in the website BirdLife.

It is then necessary to clean the dataset based the provided masks. For this task it is better to include the widest temporal range for the native breeding (from April to May), and remove duplicates (observations with the same latitude and longitude), in order to reduce computing time afterwards, as the models used do not consider frequencies.

```
months <- c(4,5,6,7)
sum1 <- db1b[db1b$month %in% months,]

pts <- sum1[,c("gbifID","decimalLatitude","decimalLongitude")]
pts <- pts[!duplicated(pts[,c("decimalLatitude","decimalLongitude")]),]
```

The resulting database (50587 observations) has been then remapped based on the grid of the climate files used later in the analysis, and again any duplicate has been removed.

```
library(ncdf4)

# Function to modify coordinates
wherenearest <- function(val,matrix) {
  dist=abs(matrix-val)
  index=which.min(dist)
  return( index )
}
```

```
# Environmental variables for the whole area, in the present
envdata <- read.table(paste(climdir,"EnvirVar/EnvirVar_NAmerica_0.txt", sep=""),
                      header=TRUE, sep=" ")
colnames(envdata)[1:2] <- c("long","lat")

#Extract latitude and longitude
lon <- as.vector(unique(envdata$long))
lat <- as.vector(unique(envdata$lat))

# modify latitude and longitude to match the grid of the climatic reconstructions
pts$decimalLatitude <- sapply(pts$decimalLatitude, function(x)
  lat[wherenearest(x,lat)])
pts$decimalLongitude <- sapply(as.numeric(pts$decimalLongitude), function(x)
  lon[wherenearest(x,lon)])

# remove duplicates
pts <- pts[!duplicated(pts[,c("decimalLatitude","decimalLongitude")]),]
```

The dataset, reduced to 4315 observations, is now ready to cleaned based on the native breeding range mask. This task takes significantly more time whit much larger datasets, this is why some of the filtering has been done beore this step.

```
library(PBSmapping)
library(rgeos)
library(maptools)

# read mask data
myspecies<-readShapePoly("Setophaga_petechia",
                        proj4string=CRS("+proj=longlat +datum=WGS84"))

sapply(slot(myspecies, "polygons"), function(x) slot(x, "ID"))

# subset it for presence=1 or 2
myspecies.sub<-myspecies[myspecies$PRESENCE<3,]
myspecies.sub<-myspecies[myspecies$ORIGIN<3,]

# select the summer component
myspecies<-myspecies.sub[myspecies.sub$SEASONAL=="2",]

# create Polysets for summer and resident component:
# merging all polygons to create a single PID, needed for later operations
myspecies<-SpatialPolygons2PolySet(myspecies)
if (length(unique(myspecies$PID))>1) {
  myspecies2<-joinPolys(myspecies,operation="UNION")
}
myspecies<-PolySet2SpatialPolygons(myspecies)

# Subset data if within polygon
# define coordinates
xy <- pts[,c(3,2)]

#transform into SpatialPointsDataFrame
df <- SpatialPointsDataFrame(coords=xy, data=pts[,1],
                             proj4string=myspecies@proj4string)
```

```
# keep only points in shapefile
pts <- df[!is.na(over(df,myspecies)),]
```

After this last filtering (leaving 3364 observations) it is possible to create the input file for the following steps. The file must have three columns, two for the geographical coordinates and a third one with a 1 for the presences (and, if needed, 0 for the absences).

```
distrib <- cbind(pts$decimalLongitude,
                 pts$decimalLatitude,
                 rep(1, length(pts$decimalLongitude)))
colnames(distrib) <- c("long","lat",ID)


# save table
write.table(distrib, file=paste(ID, "distrib.txt", sep="_"),
            quote=FALSE, sep="\t", row.names=FALSE)

# plot
png(paste(ID, "final_dataset.png", sep="_"), height=6, width=10,  units='in', res=600)
par(mfrow=c(1, 2),     # 1x2 layout
    oma=c(0, 0, 5, 0), # rows of text at the outer bottom left top right margin
    mar=c(3, 1, 3, 1), # space for row of text at ticks and to separate plots
    mgp=c(2, 1, 0))    # axis label at 2 rows distance, tick labels at 1 row

plot(newmap, col="cornsilk", bg="lightblue1", lwd=0.05, border="grey",
     xlim=c(-180,-15), ylim=c(8,90),
     main=paste("Final data\n", dim(pts)[1], "observations", sep=" "))
points(as.numeric(pts$decimalLongitude),
       as.numeric(pts$decimalLatitude),
       pch=".", col="black")

plot(newmap, col="cornsilk", bg="lightblue1", lwd=0.05, border="grey",
     xlim=c(-180,-15), ylim=c(8,90), main="Summer+resident distribution")
plot(myspecies, add=TRUE, col=col, border=NA)

title(main=paste(ID, "occurrences"),cex.main= 3,
      outer=TRUE, line=2)

dev.off()
```

#### Ecological analyses

Open the species data (if not loaded already).

```
# Species data
#distrib <- read.table("S.petechia_distrib.txt", header=TRUE, sep="\t")
```

Remove NAs from the already loaded table listing the environmental variables (no-land cells) and extract the environmental variables for the observation locations.

```
# Remove NA
envdata <-  envdata[complete.cases(envdata), ]

# Extract vars for observations
obs <- merge(distrib[,c(1,2)], envdata)
```

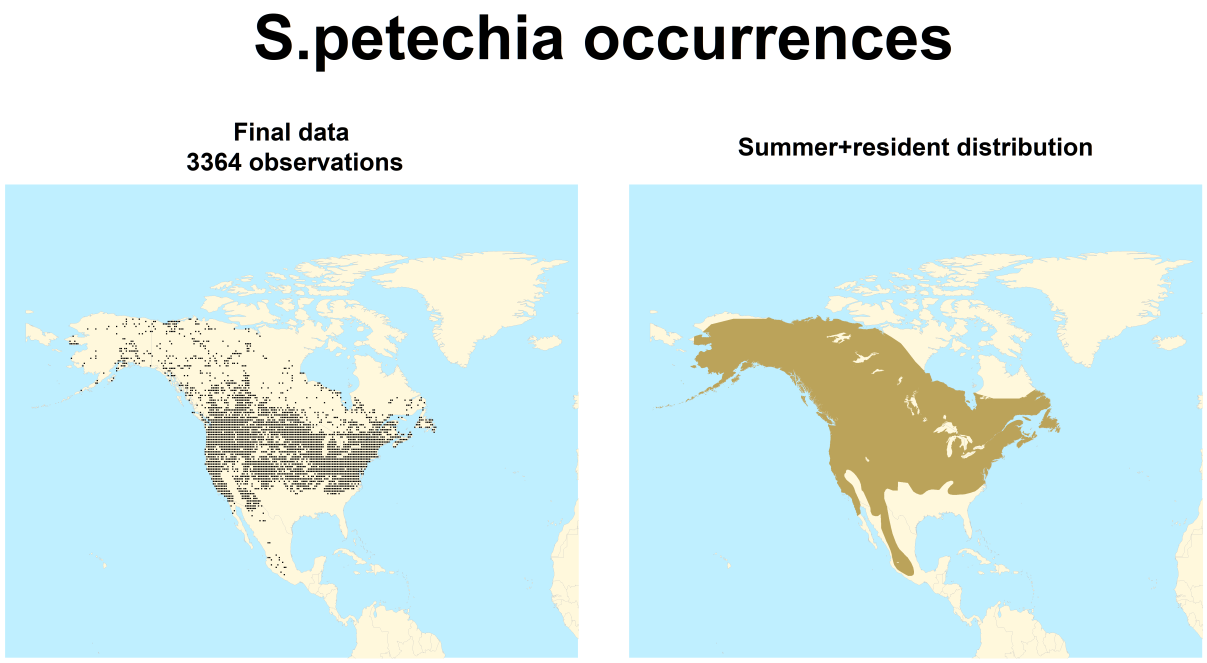


Figure 3: Final dataset

```
#head(obs)
```

Please note that the final number of observations is reduced at 3281 after removing the ones falling in no-land cells due to regridding.

##### Principal component analysis

Principal Component Analysis (PCA) based on the environmental variables.

```
# merge data in a table, last column distinguish between baseline and observations
envdata[,"set"] <- "baseline"
obs[,"set"] <- "obs"
db <- rbind(envdata,obs)

# select only climatic variables
dbMDS <- db[,vars]

# vector of colors
colors <- c(rep("black", dim(envdata)[1]),rep(col,dim(obs)[1]))

prin_comp <- prcomp(dbMDS, center=TRUE,
                    scale.=TRUE)
#extract coordinates
ind.coord <- prin_comp$x

# Eigenvalues
eig <- (prin_comp$sdev)^2
```

```r
# Variances in percentage
variance <- eig*100/sum(eig)
# Cumulative variances
cumvar <- cumsum(variance)
eig2 <- data.frame(eig=eig, variance=variance,cumvariance=cumvar)

#plot PC1 PC2
png(paste(ID, "PCA.png", sep="_"), height= 7, width=8.5,  units='in', res=600)
plot(ind.coord[,1],ind.coord[,2], col=colors, pch=20,
     ylab=paste("PC2 (",round(eig2[2,"variance"],2)," %)",sep=""),
     xlab=paste("PC1 (",round(eig2[1,"variance"],2)," %)",sep=""),
     main="PCA environmental variables")
legend("bottomright", c("Baseline","S.petechia"),pch=20,
       col=c("black",col), bty="n")
dev.off()
```

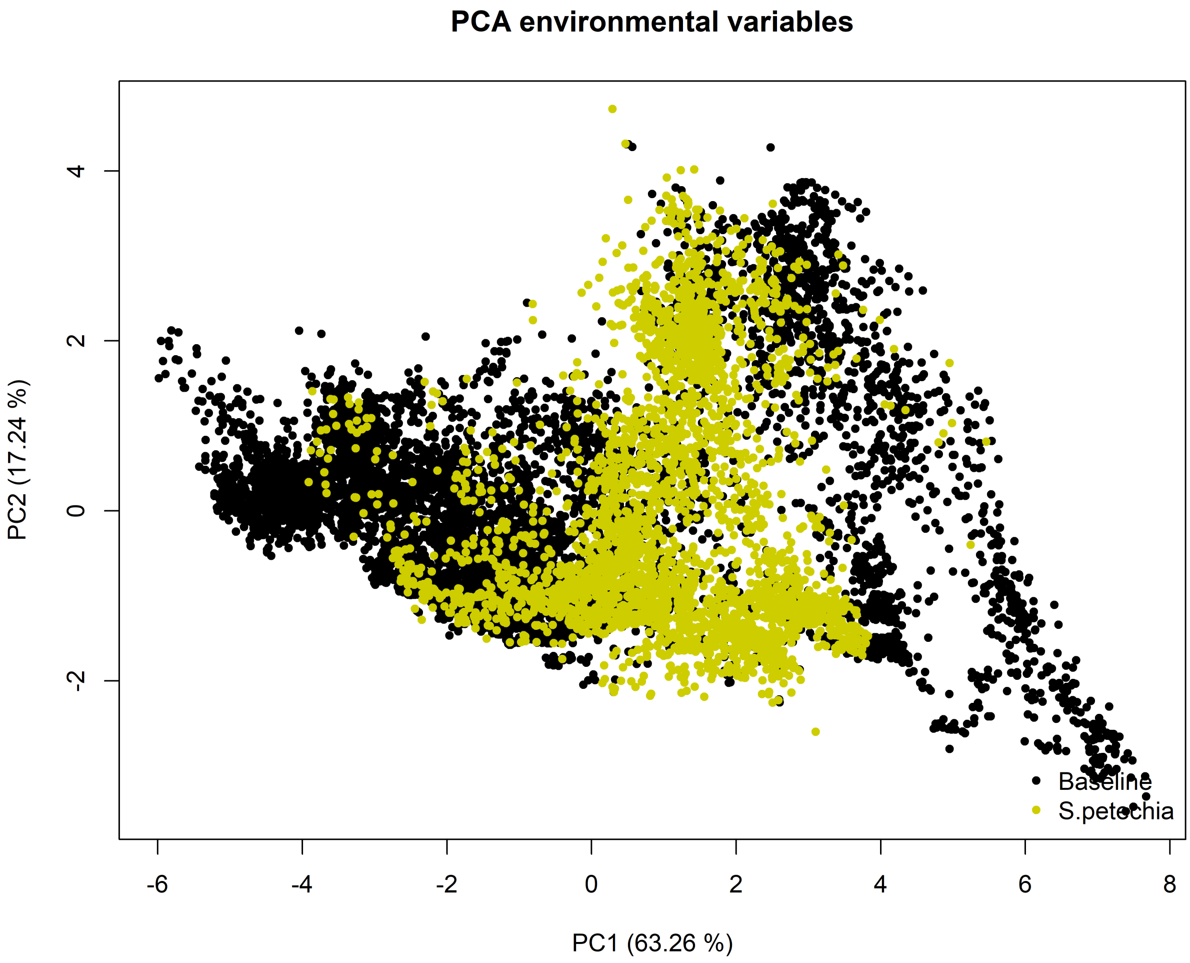


Figure 4: Principal Component Analysis (PCA) based on the environmental variables.

Plot the direction of each variable in the PCA space.

```r
#plot direction variables
var_cor_func <- function(var.loadings, comp.sdev){
  var.loadings*comp.sdev
}

# Variable correlation/coordinates
loadings <- prin_comp$rotation
sdev <- prin_comp$sdev
var.coord <- var.cor <- t(apply(loadings, 1, var_cor_func, sdev))

#head(var.coord[, 1:4])

a <- seq(0, 2*pi, length=100)

png(paste(ID, "directPCA.png", sep="_"), height=7,width=7, units='in', res=600)
plot( cos(a), sin(a), type='l', col="gray",
      xlab="PC1",  ylab="PC2")
abline(h=0, v=0, lty=2)
# Add active variables
arrows(0, 0, var.coord[, 1], var.coord[, 2],
       length=0.1, angle=15, code=2)
# Add labels tmin tmax totprec npp
text(var.coord, labels=rownames(var.coord), cex=1, adj=1, pos=3)
dev.off()
```

##### Variable distribution

Create a multiplot to compare the distribution of each variable in North America (black, on the left) with the distribution in the cells where the Yellow Warbler has been observed (on the right).

```r
library(beanplot)

# plot
png(paste(ID,"variables.png", sep="_"), height=9, width=8,  units='in', res=600)
par(mfrow=c(4, 5),
    oma=c(0, 1, 5, 0),
    mar=c(1, 1, 3, 1),
    mgp=c(0.5, 0.5, 0))

# for each environmental variable
for (x in c(3:length(vars)+2)){

  # Beanplot distributions
  beanplot(db[db$set=="baseline",x],db[db$set=="obs",x],
         bw="nrd",side="both", col=list("black",col),
         border=c("black", col), what=c(1,1,1,0),
         main=vars[x-2],xaxt='n')
}
title(main="Setophaga petechia climatic variables",cex.main= 3,
      outer=TRUE, line=2)
dev.off()
```

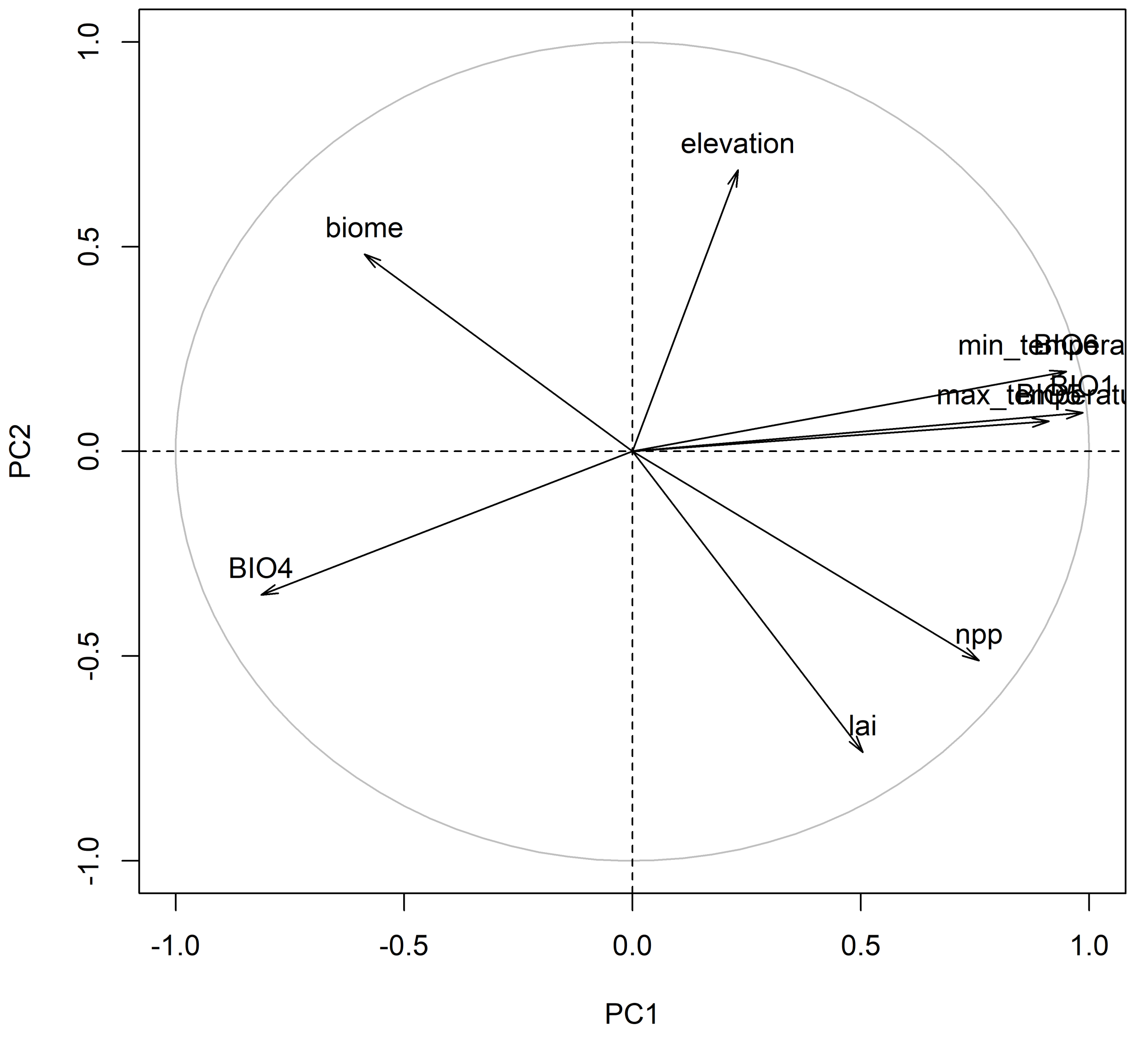


Figure 5: Plot of the direction of each variable in the PCA space.

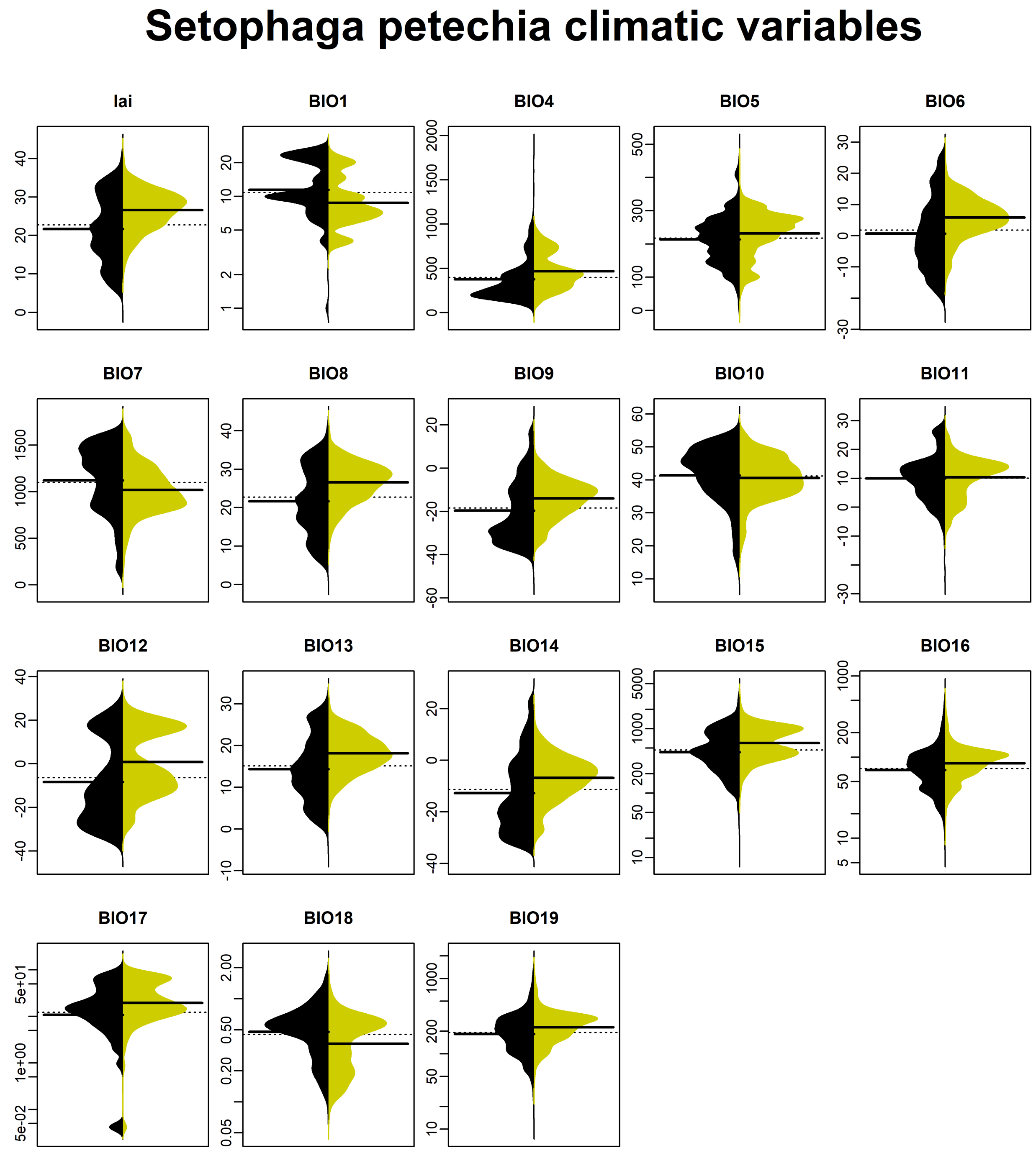


Figure 6: Comparison of the distribution of each variable in North America (black, on the left) with the distribution in the cells where the Yellow Warbler has been observed (on the right).

##### Cross-correlation

Based on the above plot and the PCA the most promising climate variables appear to be BIO1, BIO6, BIO7, BIO8, BIO9, BIO13, BIO14, BIO18, NPP and LAI. It is now necessary to calculate the cross correlation between them in order to exclude highly correlated ones.

The panel.cor function has been provided by Raquel A. Garcia, Stellenbosch University, South Africa.

```r
# function to plot (by Raquel A. Garcia, Stellenbosch University, South Africa)
panel.cor <- function(x, y, digits=2, prefix="", cex.cor)
{
  usr <- par("usr"); on.exit(par(usr))
  par(usr=c(0, 1, 0, 1))
  r <- abs(cor(x, y))
  txt <- format(c(r, 0.123456789), digits=digits)[1]
  txt <- paste(prefix, txt, sep="")
  if(missing(cex.cor)) cex <- 0.8/strwidth(txt)

  test <- cor.test(x,y)
  # borrowed from printCoefmat
  Signif <- symnum(test$p.value, corr=FALSE, na=FALSE,
                   cutpoints=c(0, 0.001, 0.01, 0.05, 0.1, 1),
                   symbols=c("***", "**", "*", ".", " "))

  text(0.5, 0.5, txt, cex=cex * r)
  text(.8, .8, Signif, cex=cex, col=2)
}

# Variables of interest
ch <-c("npp","lai","BIO1","BIO6","BIO7","BIO8","BIO9",
"BIO13","BIO14","BIO18")
# Pairwise correlation matrix
cormat <- cor(envdata[,ch])


# Plot correlation for variables of interest
png(filename=paste(ID,"correlation_vars_interest.png", sep="_"),
    width=1200, height=900)
pairs(envdata[,ch], lower.panel=panel.smooth, upper.panel=panel.cor)
dev.off()
```

In order to reduce cross-correlation we only considered variables correlated up to 0.7.

```r
library(caret)

# define highly correlated variables
hicor <- findCorrelation(cormat, cutoff=0.7)

# plot correlation for variables with correlation below 0.7
png(filename=paste(ID,"correlation_uncorr_vars.png", sep="_"),
    width=1200, height=900)
pairs(envdata[,ch[-c(hicor)]], lower.panel=panel.smooth, upper.panel=panel.cor)
dev.off()
```

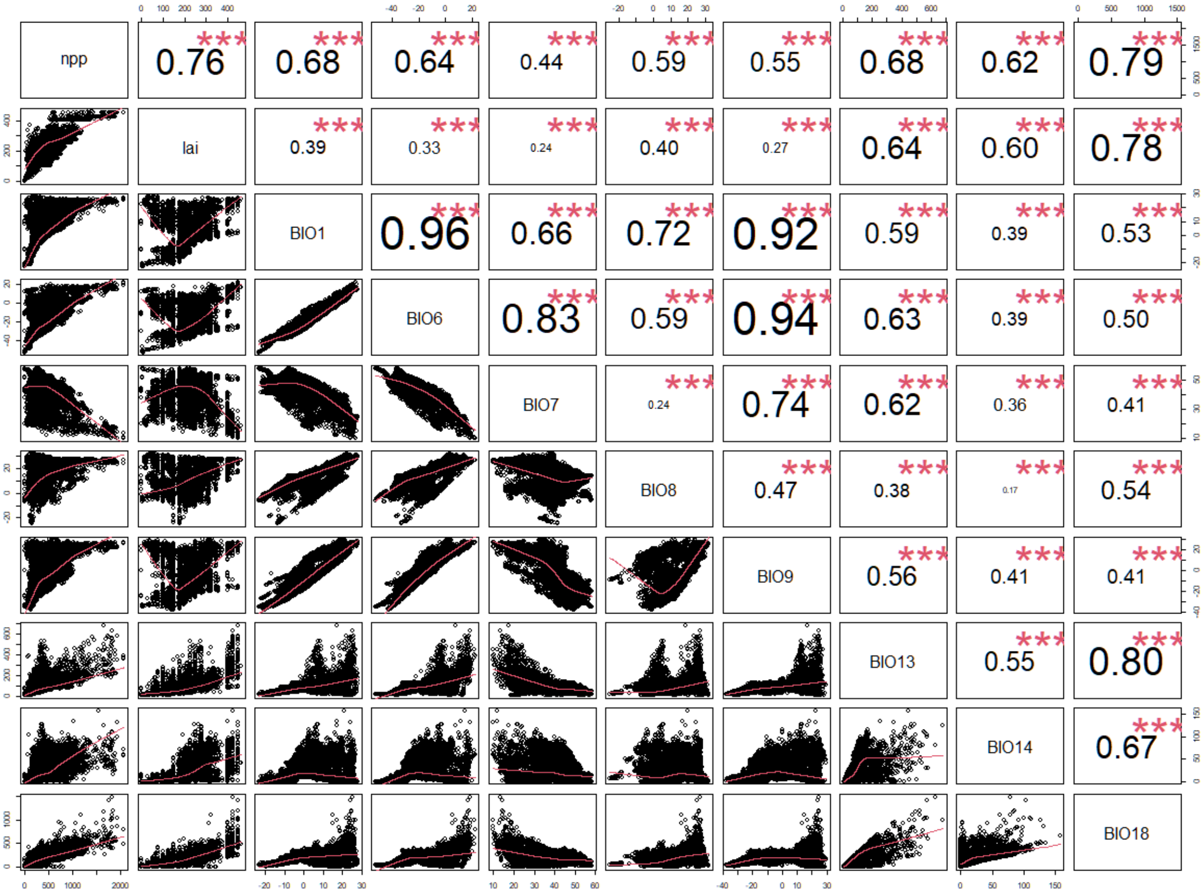


Figure 7: Correlation between all climatic variables of interest.

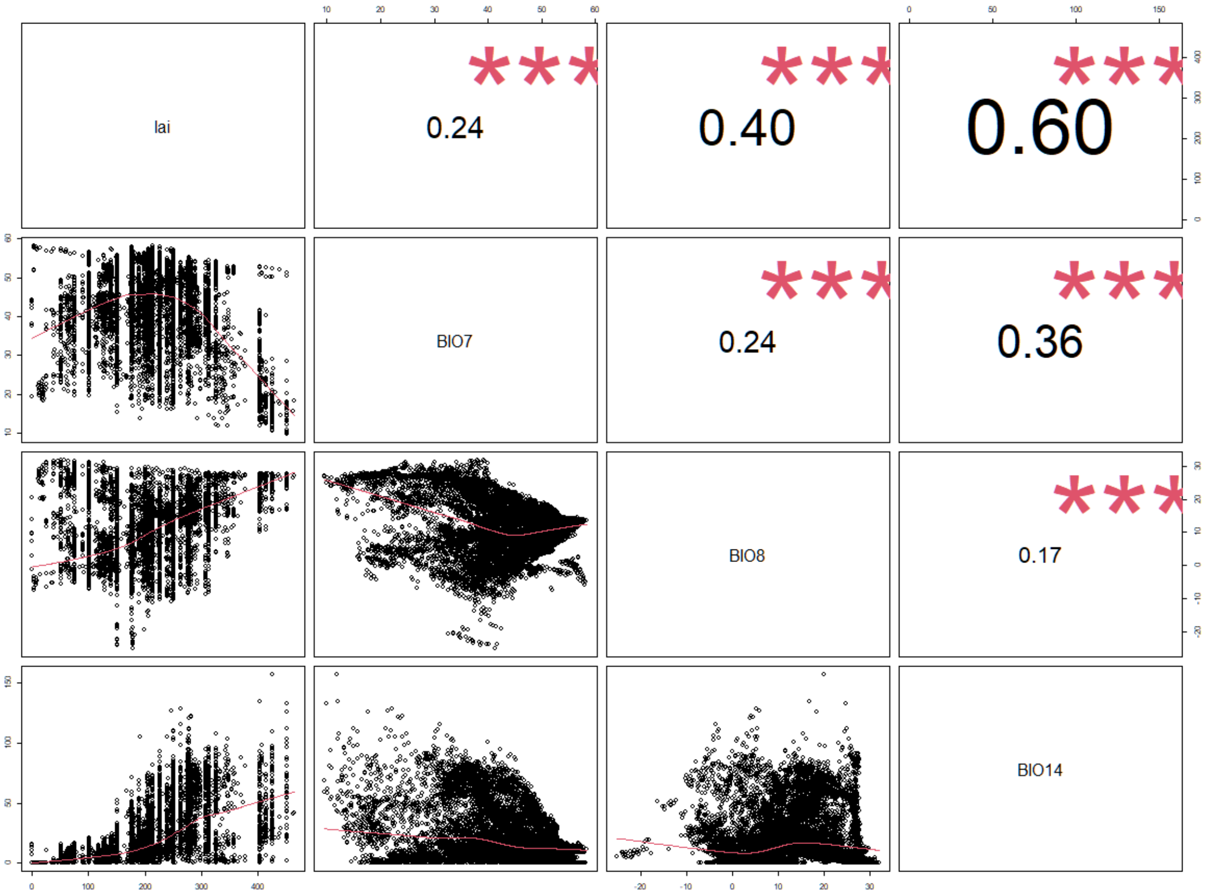


Figure 8: Correlation between chosen uncorrelated climatic variables (threshold=0.7).

#### Species distribution modelling

The following chapter details the Species Distribution modelling on the basis of the observed presences of the species, and climatic reconstructions.

Preparing the environment: load libraries, set directories for the input files and define other parameters.

```
library(RColorBrewer)

# Raster maps for climatic variables through time (NAmerica)
#basefile <- "NAmerica/"
# Directory where oputput is stored
#BiomodData <- "BiomodData/"

# uncorrelated variables of interest
vars1 <- ch[-c(hicor)]
# Color palette
#c1 <- colorRampPalette(c("khaki1","yellow3"))(6)
#c2 <- colorRampPalette(c("yellow3","black"))(8)
#cols <- c(c1[2:5],c2[2:8])
```

Please be aware that if the species name includes an underscore (e.g.*S_petechia*) it will be transformed into a point ("*S.petechia*") by the program itself when using it within the output file names.

##### Spatial thinning

In order to reduce the geographic bias associated to an uneven geographic sampling of the species we decided to thin the dataset based on a minimum distance of 70 km based on 100 repetitions with the R package SpThin.

```
library(spThin)

# Create output directory
#dir.create(file.path("Thin"))

# Spatial thinning
# t <- thin(loc.data=as.data.frame(distrib),
#           lat.col="lat", long.col="long", spec.col=ID,
#           thin.par=70, reps=100, locs.thinned.list.return=TRUE,
#           write.files=TRUE, max.files=10, out.dir="Thin/",
#           out.base="S.petechia", write.log.file=FALSE)

t_dist <- read.table("Thin/S.petechia_thin1.csv",header=TRUE,  sep=",", colClasses="numeric")
#t_dist <- read.table("Thin_100km/S.petechia_thin1.csv",header=TRUE,  sep=",")
t_dist <- t_dist[,c("long","lat",ID)]

# plot thinning
png( paste(ID,"thinned_dataset.png", sep="_"), height=6, width=13,  units='in', res=600)
par(mfrow=c(1, 2),      # 1x2 layout
    oma=c(0, 0, 5, 0), # rows of text at the outer bottom left top right margin
    mar=c(3, 1, 3, 1), # space for row of text at ticks and to separate plots
    mgp=c(2, 1, 0))    # axis label at 2 rows distance, tick labels at 1 row

plot(newmap, col="cornsilk", bg="lightblue1", border="grey", xlim=c(-180,-15), ylim=c(8,90),
     lwd=0.05, main=paste("Dataset\nN =", dim(distrib)[1], sep=" "))
points(as.numeric(distrib[,"long"]),as.numeric(distrib[,"lat"]), pch="*", col=IDcolor)
```

```
plot(newmap, col="cornsilk", bg="lightblue1", border="grey", xlim=c(-180,-15), ylim=c(8,90),
     lwd=0.05, main=paste("Thinned dataset\nN =", dim(t_dist)[1], sep=" "))
points(as.numeric(t_dist$long), as.numeric(t_dist$lat), pch="*", col="black")

title(main=paste(ID, "occurrences"),cex.main= 3,
      outer=TRUE, line=2)

dev.off()
```

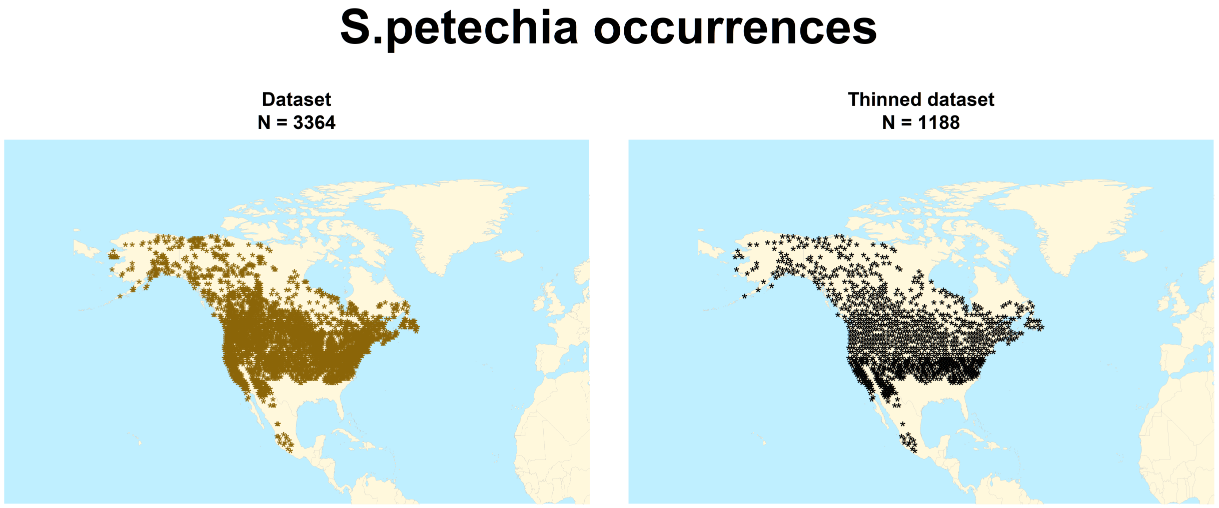


Figure 9: Original (left) and spatially thinned (right) dataset. The threshold used for spatial thinning is 70 km.

##### Geographic input file

Formatting the climate data as required for the modelling step. Loading modern day climate data (as they start with "*0_*", meaning present-day) and extracting rasters for selected variables for the formatting function.

```
library(raster)
library(rgdal)

# environmental variables for modelling
expl.var <- stack()

# create raster stack
for(v in 1:length(vars1)){
  r <- raster(paste(climdir , vars1[v], "/NAmerica_",vars1[v],"_0.grd", sep=""), RAT=FALSE)
  expl.var <- stack( expl.var, r)
}
```

##### Pseudo-absence selection

We decided to randomly draw pseudo-absences from all possible points outside the original mask, with a sample size that equals the presences. As a first step the following code identifies the points available for pseudoabsences

```r
# background data
bg <- data.frame(cbind(envdata[,c("long","lat")], S.petechia=rep(0,dim(envdata)[1])))

# define coordinates
xy <- bg[,c("long","lat")]

#transform into SpatialPointsDataFrame
df <- SpatialPointsDataFrame(coords=xy, data=bg,
                             proj4string=myspecies@proj4string)

# remove points within the mask
bg.clean <- df[is.na(over(df,myspecies)),]
bg.clean <- data.frame(bg.clean[,1:3])

# # plot
# plot(newmap, col="cornsilk", bg="lightblue1", lwd=0.05, border = "grey",
#      xlim = c(min(lon), max(lon)), ylim = c(min(lat), max(lat)))
# points(as.numeric(bg.clean[,"long"]), as.numeric(bg.clean[,"lat"]), pch="*", col="red")
```

The following code randomly drawn pseudoabsences 5 times and creates a table with information on pseudoabsences in the format of "PA.table" to be read by biomod2.

```r
# number of absences and presences
pres <- dim(t_dist)[1]
abs <- dim(t_dist)[1]

# Absences table
abs.table <- data.frame(matrix("FALSE",abs*5,7), stringsAsFactors=FALSE)
colnames(abs.table) <- c("long","lat",paste("RUN",c(1:5),sep=""))

# variable to define the first line to be written
start <- 1

for (i in 1:5){
  ix <- sample(1:dim(bg.clean)[1], abs)
  abs.table[seq(start,abs*i),1:2] <- bg.clean[ix,1:2]
  abs.table[seq(start,abs*i),2+i] <- rep("TRUE",abs)
  start <- (abs*i)+1
}
```

##### Species input file

Formatting the species distribution and extracting the relevant information as required for the modelling step. This script is based on a species input file format with three columns, two for the geographical coordinates and a third one with a 1 for the presences.

```r
#
resp.var <- as.numeric(c(rep(1, pres),rep(0,abs*5)))       # species presences + background
resp.xy <- rbind(t_dist[,c("long","lat")], abs.table[,c("long","lat")])# coordinates
resp.xy <- apply(resp.xy, 2, as.numeric)

# add presences to PA.table
pres.table <- data.frame(cbind(t_dist[,c("long","lat")]), matrix("TRUE",pres,5))
colnames(pres.table) <- c("long","lat",paste("RUN",c(1:5),sep=""))
```

```r
# merge tables for presences and pseudoabsences
PA.table <- rbind(pres.table[,-c(1,2)], abs.table[,-c(1,2)])
PA.table[] <- lapply(PA.table, as.logical)
```

Creation of the input required for biomod2 using the already defined pseudo-absences.

```r
library(biomod2)

biomodData <- BIOMOD_FormatingData(resp.var=resp.var,         # species distribution
                                   expl.var=expl.var,         # environmental variables
                                   resp.xy=resp.xy,           # coordinates of species
                                   resp.name=ID,              # name of the species column
                                   PA.strategy="user.defined",# user defined pseudo absences
                                   PA.table=PA.table,
                                   na.rm=TRUE)                # do not consider NA


png(paste(ID,"pseudoabsences_datasets.png", sep="_"), width=7, height=7, units='in', res=600)
plot(biomodData)
dev.off()
```

The resulting object can be stored on the computer executing the following code:

```r
# create BiomodData directory
dir.create("BiomodData")
# store file
save(biomodData, file=paste("./BiomodData/",ID, sep=""))
```

##### Block cross-validation

The following code uses the package blockCV to split the dataset into two parts, one for calibration and one for evaluation. It is important to reload the dataset fro the biomodData object because formating the data may remove some points (because of NA in the variables associated).

We decided to split the data into 15 vertical (North-South) columns and creating 5 different datasets in which 12 of them are used for celibration and 3 for evaluation. The output of this process is saved as *DataSplitTable* the format to be read by biomod2.

```r
library(blockCV)

# presences and absences taken from biomodData (with NA removed)
resp <- cbind(biomodData@coord,S.petechia=biomodData@data.species)
resp[] <- lapply(resp, as.numeric)

# transform into SpatialPointsDataFrame
resp <- SpatialPointsDataFrame(resp[,c("long", "lat")], resp, proj4string=crs(expl.var))

# create spatial blocks and saves plot
png(paste(ID,"spatial_blocks.png", sep="_"), height=6, width=10,  units='in', res=600)
sb <- spatialBlock(speciesData = resp,
                   species = ID,
                   rasterLayer = expl.var,
                   rows = 15,
                   k = 5,
                   selection = "systematic",
                   #iteration = 500, # find evenly dispersed folds
```

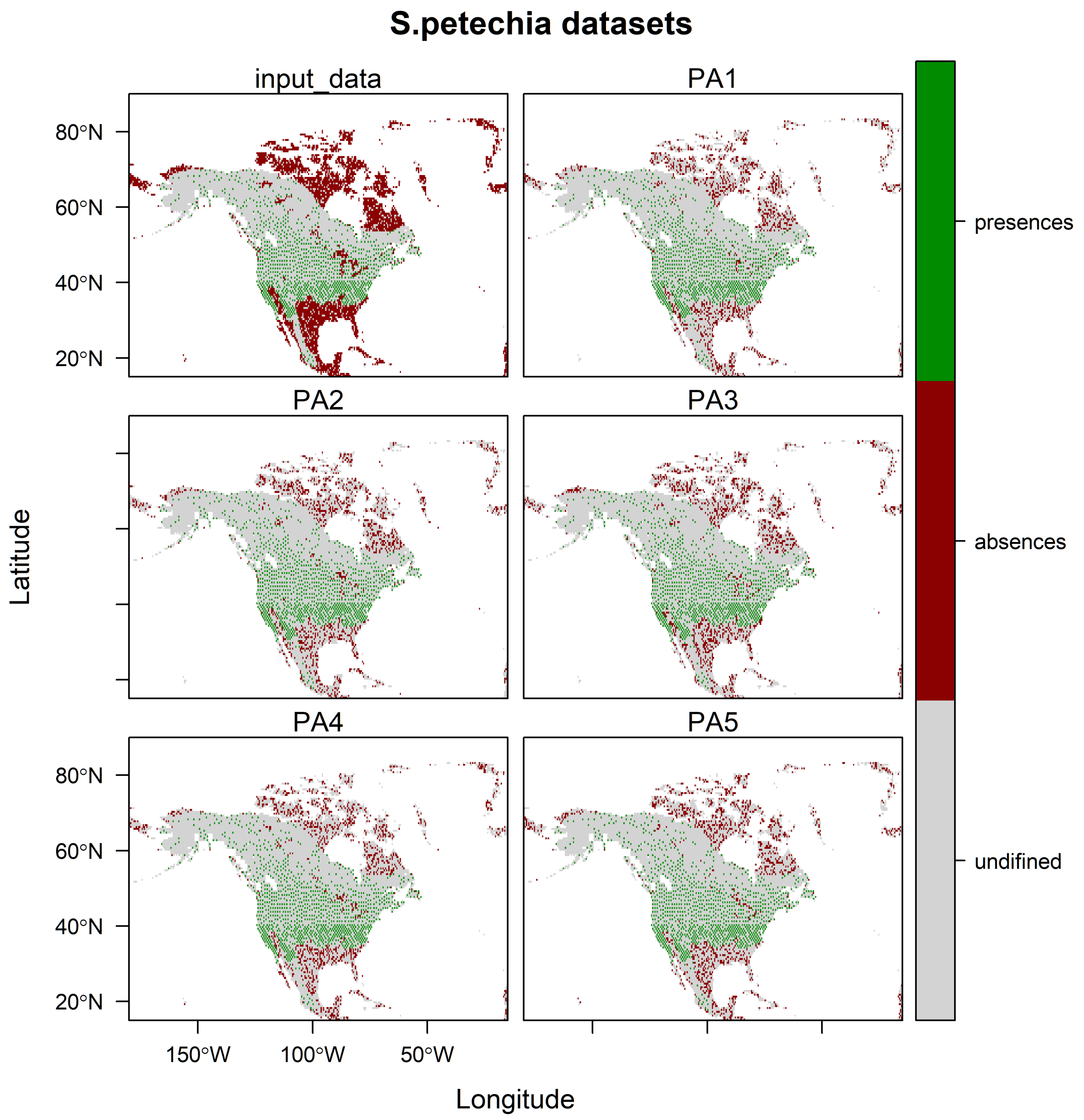


Figure 10: Pseudoabsences datasets (randomly drawn from the whole background in the same number as presences)

```
                         biomod2Format = TRUE)
dev.off()

DataSplitTable <- sb$biomodTable
```

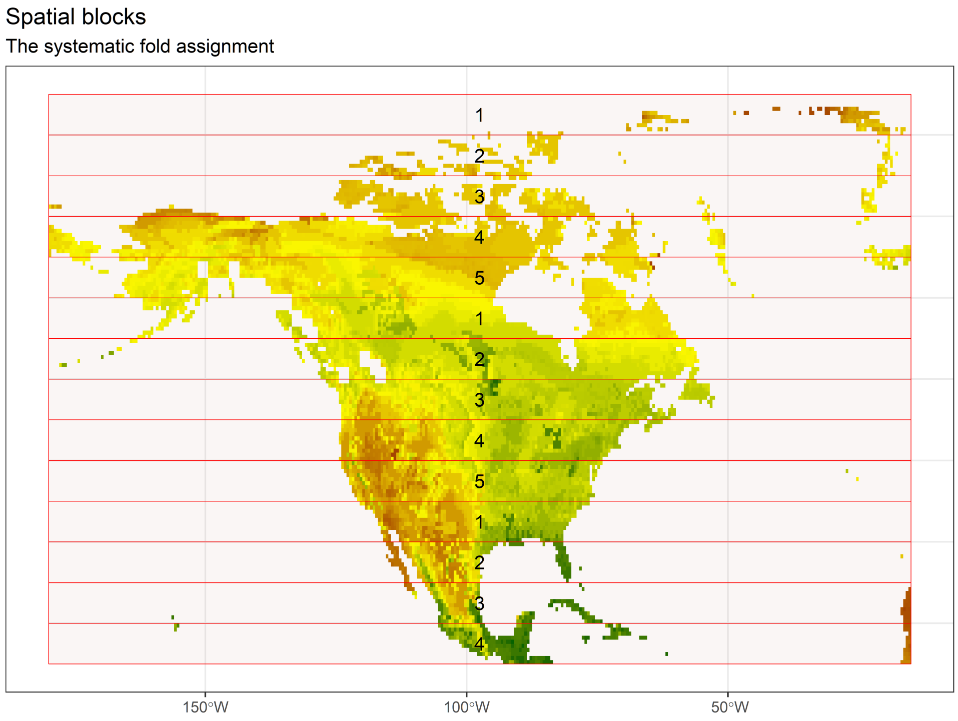


Figure 11: Spatial blocks defined by blockCV)

The following code allows to plot how the observations are used in the different runs.

```
# add coordinates to the table
dst.xy <- cbind(resp$long,resp$lat,DataSplitTable)
colnames(dst.xy)[1:2] <- c("x", "y")

# plot
png(paste(ID,"blocks_CV.png", sep="_"), width=7, height=8, units='in', res=600)
par(mfrow=c(3, 2),
    oma=c(0, 0, 5, 0),
    mar=c(1, 1, 1, 1),
    mgp=c(2, 1, 0))

for (i in c(1:5)){

  cols <- DataSplitTable[,paste("RUN",i,sep="")]
  cols[cols=="FALSE"] <- col
  cols[cols=="TRUE"] <- "black"

  plot(newmap, col="cornsilk", bg="lightblue1", lwd=0.05, border = "grey",
       xlim = c(min(lon), max(lon)), ylim = c(min(lat), max(lat)), main=paste("RUN",i,sep=""))
  points(as.numeric(dst.xy[,"x"]), as.numeric(dst.xy[,"y"]), pch="*", col=cols)
```

```
}

# title
title(main=paste(ID, "spatial blocks cross-validation"), cex.main= 2, outer=TRUE, line=3)

dev.off()
```

##### Calibrating the models

The following code allows the calibration of the models using all algorithms available in biomod2, as specified by the vector. The data is split in two parts, 80% of the data are used for calibration and 20% for evaluation based on TSS. To define the model parameters it is possible to create an object with the default options with the command *BIOMOD_ModelingOptions()*.

```
ModOptions <- BIOMOD_ModelingOptions()

mods <- c("GLM","GBM","GAM","RF") # GBM = boosted regression tree

ModelOut <- BIOMOD_Modeling(biomodData,                       # input data
                            models=mods,                      # algorithms
                            models.options=ModOptions,        # options
                            DataSplitTable=DataSplitTable,    # DataSplitTable issued by blockCV
                            #NbRunEval=3,                     # number of evaluations
                            #DataSplit=80,
                            models.eval.meth=c("TSS"),        # method for evaluat
                            SaveObj=T,
                            modeling.id=paste(ID,"_modelOut", sep=""),
                            do.full.models=FALSE,
                            rescal.all.models=T)
# model evaluations
modEval <- get_evaluations(ModelOut)

# TSS scores
write.table(modEval["TSS","Testing.data",,,], paste(ID,"models_eval.txt", sep="_"))
```

##### Projecting to the entire study area

Project the models to the entire study area (baseline period) using the same climate data used for the calibration.

```
# environmental information (same as used for calibration)
new.env <- expl.var

ProjBas <- BIOMOD_Projection(modeling.output=ModelOut, # model
                             selected.models="all",    # models to select
                             new.env= new.env,         # environmental data to project to
                             proj.name="Proj_NAmer",
                             binary.meth=c("TSS"),     # method for binary transformation
                             filtered.meth=c("TSS"),   # method to set values below zero
                             build.clamping.mask=T,    # out-of-calibration range
                             compress="xz")

# # make some plots sub-selected by str.grep argument
# png("S_petechia_proj_RF_RUN1.png", width=9, height=12, units='in', res=600)
```

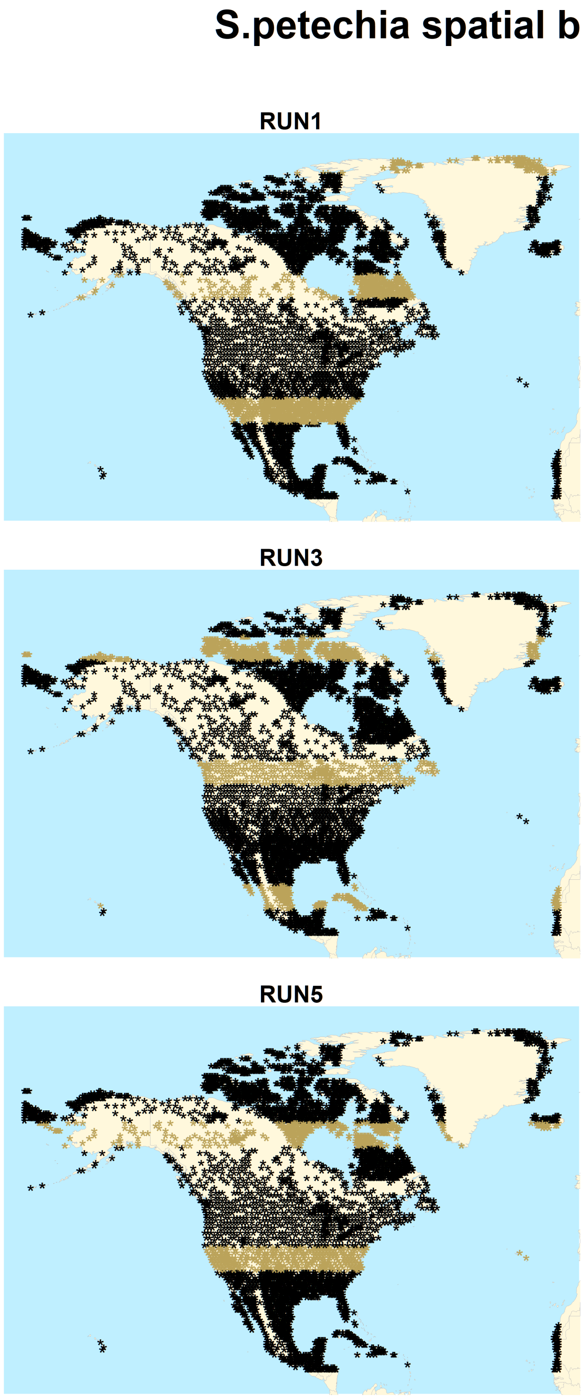


Figure 12: Spatial splitting of the data for each run: black for calibration, dark yellow for validation)

```r
# plot(ProjBas, str.grep = 'RF')
# dev.off()
```

##### Ensemble modelling

Defining the rules for generating ensembles and for evaluating them. The ensemble is built merging all algorithms together and is evaluated by TSS (threshold=0.7).

```r
EnsMod <- BIOMOD_EnsembleModeling(modeling.output=ModelOut,
                                  chosen.models="all",      # models used
                                  em.by="all",              # by algorithm
                                  #VarImport=5,              # var importance permutations
                                  eval.metric=c("TSS"),     # evaluation metric
                                  eval.metric.quality.threshold=c(0.7),
                                  models.eval.meth=c("TSS"),# evaluation method
                                  prob.mean=T,
                                  prob.median=T,
                                  committee.averaging=T,
                                  prob.mean.weight=T,
                                  prob.mean.weight.decay="proportional")
```

##### Validation of the model

Get the evaluation scores for the ensemble model

```r
eval <- get_evaluations(EnsMod)
write.table(eval, file=paste(ID,"EnsembleEvaluation.txt", sep="_"))
```

Create a table with evaluation scores, sensitivity and specificity.

```r
# change dimension names to make them more easily accessible
names(eval) <- c("mean_TSS","median_TSS","ca_TSS","wmean_TSS")

# Create a table with evaluation scores, sensitivity and specificity
out<- rbind(c(eval$mean_TSS[,1], eval$median_TSS[,1], eval$ca_TSS[,1], eval$wmean_TSS[,1]),
            c(eval$mean_TSS[,3], eval$median_TSS[,3], eval$ca_TSS[,3], eval$wmean_TSS[,3])/100,
            c(eval$mean_TSS[,4], eval$median_TSS[,4], eval$ca_TSS[,4], eval$wmean_TSS[,4])/100)
```

Plot in a single figure three barplots showing the evaluation scores based on TSS for each of the five independent evaluations.

```r
# index for the maximum TSS
ix <- which.max(out[1,])

# Plot barplots of evaluation
colnames(out) <- c("Mean","Median","Committee\naverage","Weighted\nmean")
rownames(out) <- c("Evaluation score", "Sensitivity","Specificity")

png(paste(ID,"evaluation_scores_ensemble.png", sep="_"), width=9, height=6, units='in', res=600)
layout(matrix(c(1,2), ncol=1, byrow=TRUE), heights=c(3.8,0.7))
par(oma=c(0, 0, 4, 0), # rows of text at the outer bottom left top right margin
    mar=c(2, 3, 3, 1))#, # space for row of text at ticks and to separate plots

barplot(t(out), beside=TRUE, col=cS[c(6,2,8,4)], border=NA, ylim=c(0,1))

par(mai=c(0,0,0,0))
```

```
plot.new()
legend("center", colnames(out), fill=cS[c(6,2,8,4)], border=NA, bty="n",ncol=4)

title(main=paste("Evaluation score (TSS) -", ID, sep=" "), cex.main=1.5, outer=TRUE, line=1)

dev.off()
```

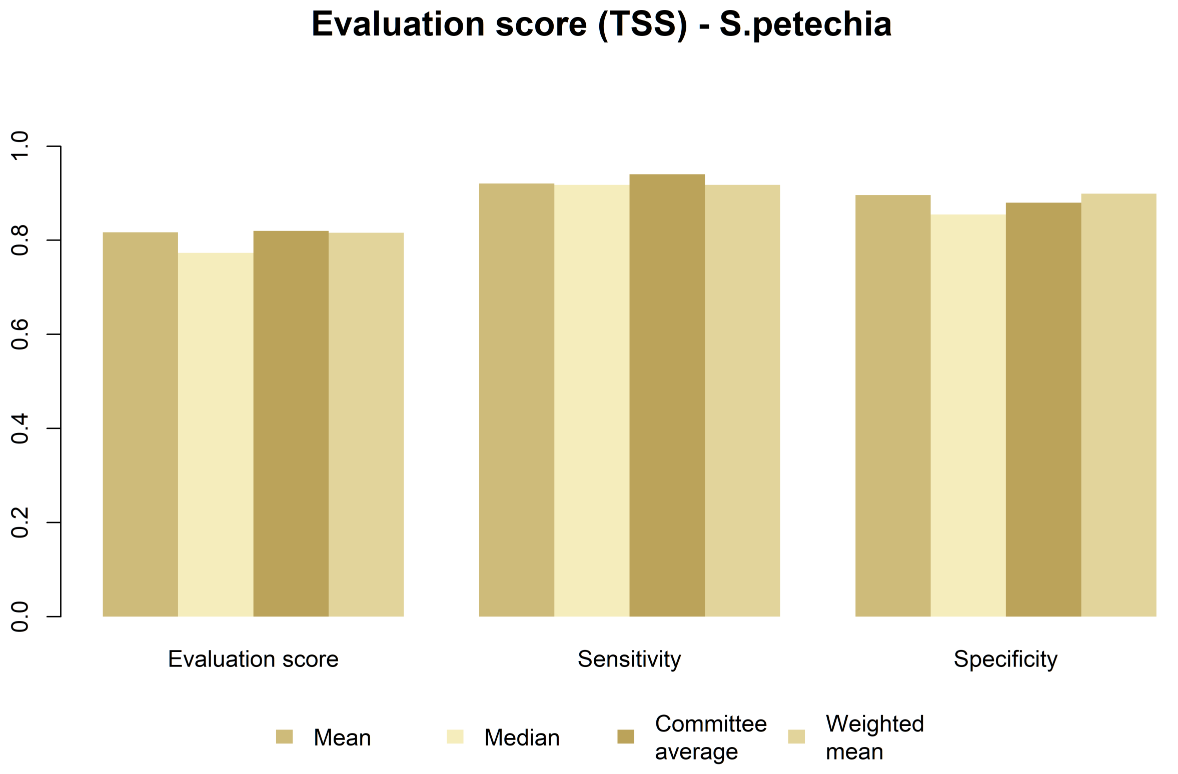


Figure 13: Ensemble model evaluation

##### Ensemble forecasting

The following code allows projecting the ensemble to build the consensus projections based on the same climatic variables already used for the other steps (new.env, which is a copy of expl.var)

```
EnsBas <- BIOMOD_EnsembleForecasting(EM.output=EnsMod,          # output from ensemble modelling
                                     projection.output=ProjBas, # output from projection
                                     selected.models="all",     # model chosen
                                     binary.meth=c("TSS"),      # method for binary transf
                                     filtered.meth=c("TSS"),    # method for filtering
                                     compress=T)                # compression
```

Exploring the output: loading the binary consensus projections:

```
#list of files in directory
filelist <- list.files(paste(ID,"/proj_Proj_NAmer/", sep=""))
# selecting consensus projections
enslist <- filelist[grep("ensemble", filelist)]
```

```
enspbin <- stack(paste(ID,"/proj_Proj_NAmer/", enslist[2], sep=""))
```

and plotting them:

```
ens.methods <- c("Mean", "Median", "Committee average", "Weighted mean")

# Read ice mask
r <- raster(paste(icedir, "Mask_0.nc", sep=""))

png(paste(ID,"ensemble_projection.png", sep="_"),
    width=12, height=9, units='in', res=600)
par(mfrow=c(2, 2),
    oma=c(0, 0, 0, 1),
    mar=c(3, 3, 2, 5),
    mgp=c(2, 1, 0))

for (x in 1:4) {
  #plot selected rasters (ensemble models)
  plot(raster(enspbin,c(x))/1000, main=paste("Modern projection", ens.methods[x], sep=" - "),
       axes=FALSE, box=FALSE, colNA="lightblue1", col=rasterColors, breaks=seq(0, 1, 0.25),)

  # plot observation points
  #points(resp.xy[,"long"], resp.xy[,"lat"], pch=".", cex=0.5)

  # plot ice
  plot(r, add=TRUE, col=c(NA,"honeydew3"), border=NA, useRaster=F, legend=FALSE)
}

dev.off()
```

#### Projection backwards in time

The following code repeat the whole process in order to project the potential distribution of the species backwards in time.

```
dir.create(file.path("Past"), showWarnings=FALSE)

# list of kyrs ago available
ref <- c(1:22, seq(24,50,2))

# for each kyrs
for (i in ref){
  # Read ice mask
  ice <- raster(paste(icedir, "Mask_", i, ".nc", sep=""))

  # create raster stack
  past.env <- stack()

  # loading climate data for selected variables
  for(v in 1:length(vars1)){
  r <- raster(paste(climdir, vars1[v], "/NAmerica_",vars1[v],"_",i, ".grd", sep=""),
              RAT=FALSE)
  past.env <- stack(past.env, r)
  }
  names(past.env)<- vars1
```

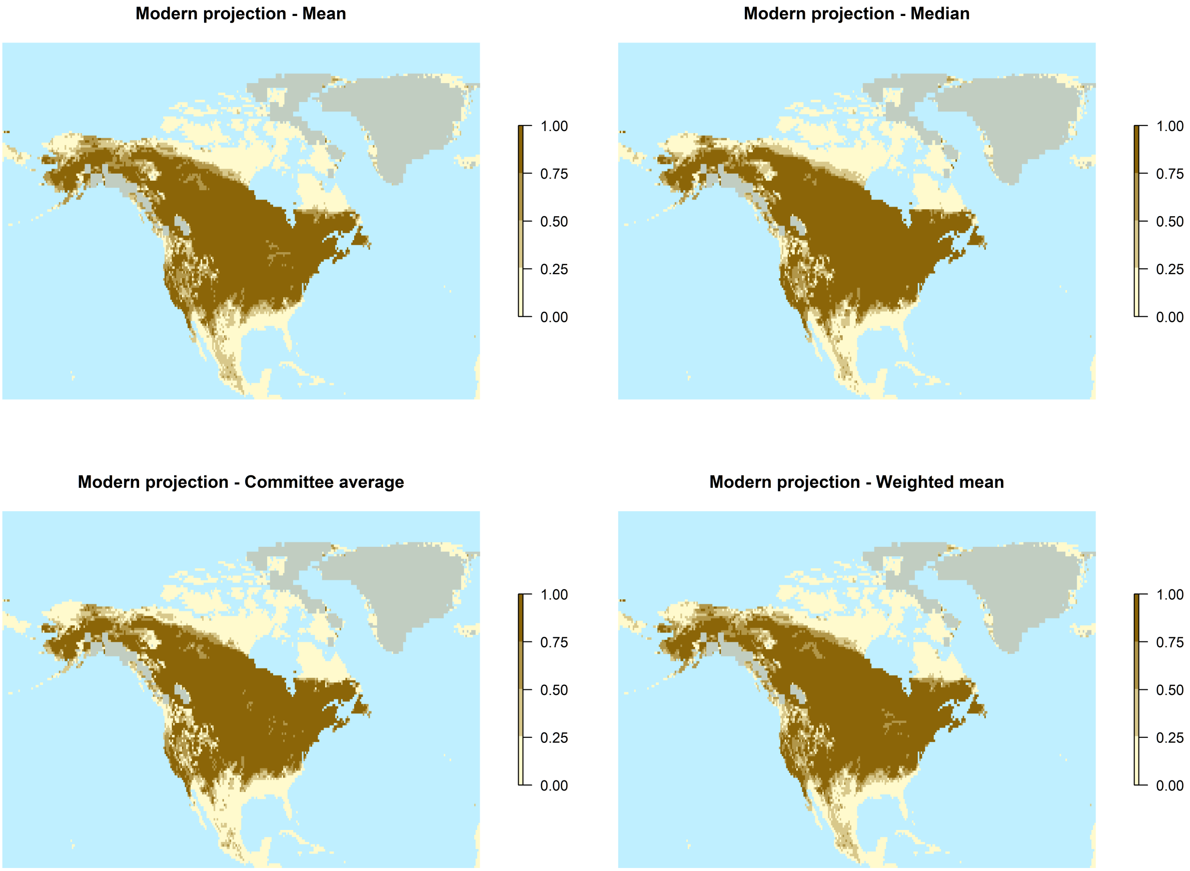


Figure 14: Plot of the ensemble projections

```
  # Project model into the past
  ProjPast <- BIOMOD_Projection(modeling.output=ModelOut,  # model
                              selected.models="all",        # model to select
                              new.env= past.env,           # environmental data to project to
                              proj.name=paste(i,"_NAmer", sep=""),
                              binary.meth="TSS",           # method for binary transformation
                              filtered.meth="TSS",         # method to set values below zero
                              build.clamping.mask=T,       # out-of-calibration range
                              compress="xz")

  # Project ensemble to the past
  EnsPast <- BIOMOD_EnsembleForecasting(EM.output=EnsMod,           # output from ensemble mod
                                      projection.output=ProjPast,  # output from projection
                                      selected.models="all",       # model chosen
                                      binary.meth=c("TSS"),        # eval method for bin transf
                                      filtered.meth=c("TSS"),      # eval method for filtering
                                      compress=T)                  # compression?


  filelist <- list.files(paste(ID,"/proj_",i,"_NAmer/", sep="")) #list of files in directory
  enslist <- filelist[grep("ensemble", filelist)]  # selecting consensus projections
  enspbin <- stack(paste(ID,"/proj_",i,"_NAmer/", enslist[2], sep=""))

  png(paste("Past/",ID,"_",i,"_all_ensemble_projections.png", sep=""),
      width=12, height=9, units='in', res=600)

  op<-par(mfrow=c(2, 2),
          oma=c(0, 0, 0, 1),
          mar=c(3, 3, 2, 5),
          mgp=c(2, 1, 0))

  for (x in 1:4) {
    # plot selected rasters (ensemble models)
    plot(raster(enspbin,c(x)/1000), main=paste("Projection",i,"k years ago -", ens.methods[x],
                                               sep=" "),
         axes=FALSE, box=FALSE, colNA="lightblue1", col=rasterColors, breaks=seq(0, 1, 0.25),)
    # add ice
    plot(ice, add=TRUE, col=c(NA,"honeydew3"), border=NA, useRaster=F, legend=FALSE)
    }
  dev.off()


  for (x in 1:4) {
    png(paste("Past/", ID, "_", i, "_", ens.methods[x],"_ensemble_projection.png", sep=""),
        width=12, height=9, units='in', res=600)
    plot(raster(enspbin,c(x)/1000), main=paste(ens.methods[x], i, "k years ago", sep=" "),
         col=rasterColors, breaks=seq(0, 1, 0.25),)
    plot(ice, add=TRUE, col=c(NA,"honeydew3"), border=NA, useRaster=F, legend=FALSE)
    dev.off()
  }

}
```

The following code creates a plot showing the mask, the data and the projection of the ensemble in four key periods: present-day, middle Holocene (6kyrs ago), Last Glacial Maximum (21 kyrs ago) and 50 kyrs ago

```
# read raster modern
filelist <- list.files("S.petechia/proj_Proj_NAmer/") #list of files in directory
enslist <- filelist[grep("ensemble", filelist)]  # selecting consensus projections
enspbin <- stack(paste("S.petechia/proj_Proj_NAmer/", enslist[2], sep=""))
pr <- raster(enspbin,3)/1000

# Holocene
filelist <- list.files("S.petechia/proj_6_NAmer/")
enslist <- filelist[grep("ensemble", filelist)]
enspbin <- stack(paste("S.petechia/proj_6_NAmer/", enslist[2], sep=""))
HOL <- raster(enspbin,3)/1000

# LGM
filelist <- list.files("S.petechia/proj_21_NAmer/")
enslist <- filelist[grep("ensemble", filelist)]
enspbin <- stack(paste("S.petechia/proj_21_NAmer/", enslist[2], sep=""))
LGM <- raster(enspbin,3)/1000

# 50k
filelist <- list.files("S.petechia/proj_50_NAmer/")
enslist <- filelist[grep("ensemble", filelist)]
enspbin <- stack(paste("S.petechia/proj_50_NAmer/", enslist[2], sep=""))
k50 <- raster(enspbin,3)/1000

# Ice
r <- raster(paste(icedir,"Mask_0.nc", sep=""))
r12 <- raster(paste(icedir,"Mask_12.nc", sep=""))
r21 <- raster(paste(icedir,"Mask_21.nc", sep=""))
r50 <- raster(paste(icedir,"Mask_50.nc", sep=""))

# plot
png(paste(ID, "6_plot.png",sep="_"), height=4.5, width=9,  units = 'in', res = 600)
par(mfrow = c(2, 3),      # 2x2 layout
    oma = c(0, 0, 5, 2), # rows of text at the outer bottom left top right margin
    mar = c(1, 2, 1, 3), # space for row of text at ticks and to separate plots
    mgp = c(2, 1, 0))     # axis label at 2 rows distance, tick labels at 1 row

# occurrences
plot(newmap, col="cornsilk", bg="lightblue1", lwd=0.05, border = "grey",
     xlim = c(min(lon), max(lon)), ylim = c(min(lat), max(lat)), main="Thinned observations")
points(as.numeric(t_dist[,"long"]), as.numeric(t_dist[,"lat"]), pch="*", col="black")

# mask
plot(newmap, col="cornsilk", bg="lightblue1", lwd=0.05, border = "grey",
     xlim = c(min(lon), max(lon)), ylim = c(min(lat), max(lat)),
     main="Summer distribution")
plot(myspecies, add=TRUE, col=col, border=NA)


# Ensemble projection modern day
plot(pr,main="Present day", axes=FALSE, box=FALSE, colNA="lightblue1",
```

```r
     col=rasterColors, breaks=seq(0, 1, 0.25))
plot(r, add=TRUE, col=c(NA,"honeydew3"), border=NA, useRaster=F, legend=FALSE)

# Ensemble projection Holocene
plot(HOL,main="Early Holocene", axes=FALSE, box=FALSE, colNA="lightblue1",
     col=rasterColors, breaks=seq(0, 1, 0.25))
plot(r12, add=TRUE, col=c(NA,"honeydew3"), border=NA, useRaster=F, legend=FALSE)

# Ensemble projection LGM
plot(LGM,main="LGM", axes=FALSE, box=FALSE, colNA="lightblue1",
     col=rasterColors, breaks=seq(0, 1, 0.25))
plot(r21, add=TRUE, col=c(NA,"honeydew3"), border=NA, useRaster=F, legend=FALSE)

# Ensemble projection past
plot(k50,main="50 kya", axes=FALSE, box=FALSE, colNA="lightblue1",
     col=rasterColors, breaks=seq(0, 1, 0.25))
plot(r50, add=TRUE, col=c(NA,"honeydew3"), border=NA, useRaster=F, legend=FALSE)

# title
title(main=paste("S. petechia (N=",dim(t_dist)[1],")",sep=""),cex.main= 2,outer = TRUE,
      line = 2)

dev.off()


# plot
png(paste(ID, "2_plot.png",sep="_"), height=4, width=9,  units = 'in', res = 600)
par(mfrow = c(1, 2),     # 2x2 layout
    oma = c(0, 0, 5, 0), # rows of text at the outer bottom left top right margin
    mar = c(1, 1, 1, 1), # space for row of text at ticks and to separate plots
    mgp = c(1, 1, 0))    # axis label at 2 rows distance, tick labels at 1 row

# mask
plot(newmap, col="cornsilk", bg="lightblue1", lwd=0.05, border = "grey",
     xlim = c(min(lon), max(lon)), ylim = c(min(lat), max(lat)),
     main="Observed")
plot(myspecies, add=TRUE, col=rasterColors[4], border=NA)


# Ensemble projection modern day
plot(pr,main="Reconstructed with SDMs", axes=FALSE, box=FALSE, colNA="lightblue1",
     col=rasterColors[c(1,4)], breaks=seq(0, 1, 0.5), legend=FALSE)
plot(r, add=TRUE, col=c(NA,"honeydew3"), border=NA, useRaster=F, legend=FALSE)

# title
title(main="Summer distribution S. petechia",cex.main= 2,outer = TRUE, line = 2)

dev.off()



# gradient plot
png(paste(ID, "2_plot_gradient.png",sep="_"), height=4, width=9,  units = 'in', res = 600)
```

```r
par(mfrow = c(1, 2),      # 2x2 layout
    oma = c(0, 0, 5, 1), # rows of text at the outer bottom left top right margin
    mar = c(1, 1, 1, 1), # space for row of text at ticks and to separate plots
    mgp = c(1, 1, 0))     # axis label at 2 rows distance, tick labels at 1 row

# mask
plot(newmap, col="cornsilk", bg="lightblue1", lwd=0.05, border = "grey",
     xlim = c(min(lon), max(lon)), ylim = c(min(lat), max(lat)),
     main="Geographic range")
plot(myspecies, add=TRUE, col=rasterColors[4], border=NA)


# Ensemble projection modern day
plot(pr,main="Estimated potential distribution", axes=FALSE, box=FALSE, colNA="lightblue1",
     col=rasterColors, breaks=seq(0, 1, 0.25))
plot(r, add=TRUE, col=c(NA,"honeydew3"), border=NA, useRaster=F, legend=FALSE)

dev.off()
```

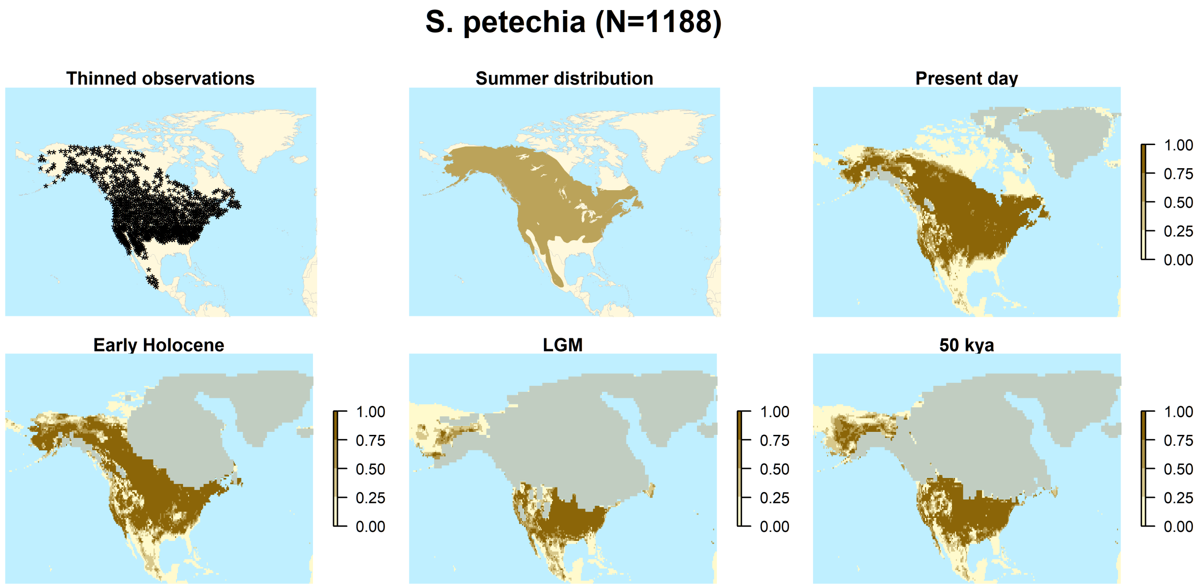


Figure 15: Final plot: original dataset, thinned dataset, summer distribution, and projection in 4 different periods of time.

#### Plot distribution through time

The following code plots the potential distribution of the Yellow Warbler in six time slices of interest: 46, 21, 13, 11, 9, and 5 ky ago.

```r
# plot
png(paste(ID, "6_distrib.png",sep="_"), height=8, width=7, units = 'in', res = 600)
par(mfrow = c(3, 2),      # 2x2 layout
    oma = c(0, 0, 5, 2), # rows of text at the outer bottom left top right margin
    mar = c(1, 2, 1, 3), # space for row of text at ticks and to separate plots
    mgp = c(2, 1, 0))     # axis label at 2 rows distance, tick labels at 1 row
```

```r
for (i in c(46,21,13,11,9,5)){
  # Read ice mask
  ice <- raster(paste(icedir, "Mask_", i, ".nc", sep=""))

  filelist <- list.files(paste("S.petechia/proj_", i, "_NAmer/", sep=""))
  enslist <- filelist[grep("ensemble", filelist)]
  enspbin <- stack(paste("S.petechia/proj_",i,"_NAmer/", enslist[2], sep=""))
  d <- raster(enspbin,3)/1000

  plot(d, main=paste(i,"kya"), axes=FALSE, box=FALSE, colNA="lightblue1",
     col=rasterColors, breaks=seq(0, 1, 0.25))
  plot(ice, add=TRUE, col=c(NA,"honeydew3"), border=NA, useRaster=F, legend=FALSE)

}

# title
title(main="S. petechia, potential range through time" ,cex.main= 2,outer = TRUE,
      line = 2)

dev.off()



for (i in c(46,21,13,11,9,5)){
  # Read ice mask
  ice <- raster(paste(icedir, "Mask_", i, ".nc", sep=""))
  for (kk in c(1:length(vars1))){
  d <- raster(paste(climdir, vars1[kk], "/NAmerica_",vars1[kk],"_",i, ".grd", sep=""), RAT=FALSE)

  png(paste(ID, "_", vars1[kk], "_", i, "kya_plot.png",sep=""), height=4.5, width=5,
      units='in', res=600)
  plot(d, main=paste(vars1[kk], i,"kya"), axes=FALSE, box=FALSE, colNA="#B9E6FF")
  plot(ice, add=TRUE, col=c(NA,"honeydew3"), border=NA, useRaster=F, legend=FALSE)
dev.off()

  }
}
```

##### Save output(s) as netcdf file

The following code saves the projections through time as netcdf files.

```r
# read file for the present and create rasters for stats of interest
filelist <- list.files(paste(ID,"/proj_Proj_NAmer/", sep="")) #list of files in directory
enslist <- filelist[grep("ensemble", filelist)]  # selecting consensus projections
enspbin <- stack(paste(ID,"/proj_Proj_NAmer/", enslist[2], sep=""))

# create raster for each statistic
mea <- raster(enspbin,c(1))
med <- raster(enspbin,c(2))
ca <- raster(enspbin,c(3))
wm <- raster(enspbin,c(4))
```

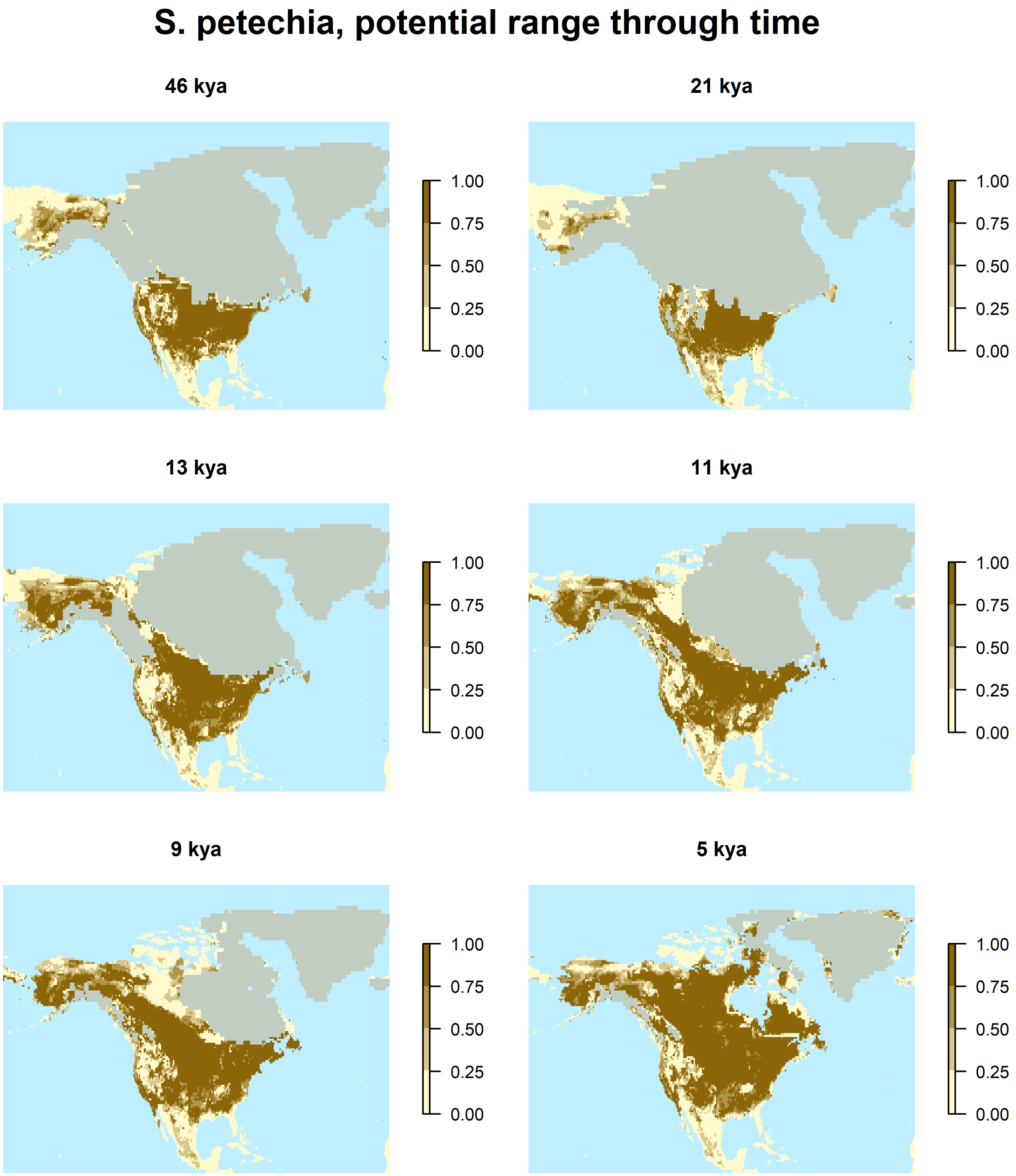


Figure 16: Projection of the potential range of the Yellow Warbler in 6 different periods of time.

```r
# do the same for all periods, stack rasters and save as netcdf file
for (i in ref){

  filelist <- list.files(paste(ID,"/proj_",i,"_NAmer/", sep="")) #list of files in directory
  enslist <- filelist[grep("ensemble", filelist)]  # selecting consensus projections
  enspbin <- stack(paste(ID,"/proj_",i,"_NAmer/", enslist[2], sep=""))

  mea <- stack(mea, raster(enspbin,c(1)))
  med <- stack(med, raster(enspbin,c(2)))
  ca <- stack(ca, raster(enspbin,c(3)))
  wm <- stack(wm, raster(enspbin,c(4)))
}

  writeRaster(mea, paste("Yellow_warbler_",ens.methods[1],".nc", sep=""),
              overwrite=TRUE, format="CDF", varname=ens.methods[1],
              longname=paste("Probability (scale: 0-1000) of Yellow warbler presence
                             from SDM analyses", ens.methods[1], sep=" - "),
              xname="Longitude", yname="Latitude", zname="Time (kyrs ago)")

  writeRaster(med, paste("Yellow_warbler_",ens.methods[2],".nc", sep=""),
              overwrite=TRUE, format="CDF", varname=ens.methods[2],
              longname=paste("Probability (scale: 0-1000) of Yellow warbler presence
                             from SDM analyses", ens.methods[2], sep=" - "),
              xname="Longitude", yname="Latitude", zname="Time (kyrs ago)")

  writeRaster(ca, paste("Yellow_warbler_",ens.methods[3],".nc", sep=""),
              overwrite=TRUE, format="CDF", varname=ens.methods[3],
              longname=paste("Probability (scale: 0-1000) of Yellow warbler presence
                             from SDM analyses", ens.methods[3], sep=" - "),
              xname="Longitude", yname="Latitude", zname="Time (kyrs ago)")

  writeRaster(wm, paste("Yellow_warbler_",ens.methods[4],".nc", sep=""),
              overwrite=TRUE, format="CDF", varname=ens.methods[4],
              longname=paste("Probability (scale: 0-1000) of Yellow warbler presence
                             from SDM analyses", ens.methods[4], sep=" - "),
              xname="Longitude", yname="Latitude", zname="Time (kyrs ago)")


#ncin <- nc_open("Yellow_warbler_Mean.nc")
#ncin
#nc_close(ncin)
```
